# Asymmetric histone inheritance regulates stem cell fate in *Drosophila* midgut

**DOI:** 10.1101/2020.08.15.252403

**Authors:** Emily Zion, Xin Chen

## Abstract

A fundamental question in developmental biology is how distinct cell fates are established and maintained through epigenetic mechanisms in multicellular organisms. Here, we report that preexisting (old) and newly synthesized (new) histones H3 and H4 are asymmetrically inherited by the distinct daughter cells during asymmetric division of *Drosophila* intestinal stem cells (ISCs). By contrast, in symmetrically dividing ISCs that produce two self-renewed stem cells, old and new H3 and H4 show symmetric inheritance patterns. These results indicate that asymmetric histone inheritance is tightly associated with the distinct daughter cell fates. To further understand the biological significance of this asymmetry, we express a mutant histone that compromises asymmetric histone inheritance pattern. We find increased symmetric ISC division and ISC tumors during aging under this condition. Together, our results demonstrate that asymmetric histone inheritance is important for establishing distinct cell identities in a somatic stem cell lineage, consistent with previous findings in asymmetrically dividing male germline stem cells in *Drosophila*. Therefore, this work sheds light on the principles of histone inheritance in regulating stem cell fate *in vivo*.

## Introduction

In multicellular organisms, tissue homeostasis could be achieved by asymmetric cell division (ACD) of adult stem cells, which produces a self-renewed stem cell and a differentiating daughter cell. This division allows the stem cell pool to be retained while giving rise to differentiating cells that replace lost cells due to turnover or tissue damage. ACD is also involved in tissue regeneration, serving as an important mechanism in maintaining the proper physiological functions of the corresponding tissue or organ (Kahney et al., 2017; Knoblich, 2008; Morrison and Spradling, 2008; Venkei and Yamashita, 2018). Disruption of this precisely regulated cell division can result in the dysregulation of stem cells, leading to cancer or tissue degeneration (Clevers, 2005; Knoblich, 2010; Morrison and Kimble, 2006).

The faithful establishment of cellular identities involves expressing a specific subset of genes and silencing other genes in a given cell type, allowing for the cell to adopt a particular morphology and execute specialized functions. Epigenetic mechanisms, such as DNA methylation and histone modifications, are heritable changes that affect gene expression without altering the DNA sequences. These mechanisms enable different cell types within a multicellular organism to establish distinct cellular identities while maintaining the same genetic information. Canonical histone proteins H3, H4, H2A, and H2B are incorporated into DNA as an octamer structure, forming the fundamental units of chromatin. It is well known that chromatin structure affects gene expression; however, it remains largely unclear how epigenetic information is maintained or changed during cell divisions *in vivo* to produce cells with distinct identities in multicellular organisms (Ahmad and Henikoff, 2018; Allis and Jenuwein, 2016; Badeaux and Shi, 2013; Kouzarides, 2007; Yadav et al., 2018; Young et al., 2010).

Previous studies have shown that H3 and H4 histones can be inherited asymmetrically, with preexisting (old) histones retained in the self-renewed stem cell while newly synthesized (new) histones enriched in the differentiating daughter cell during ACD of the *Drosophila* male germline stem cells (GSCs) (Tran et al., 2012; Wooten et al., 2019). In contrast, old and new H2A and H2B are inherited symmetrically during ACD of male GSCs (Wooten et al., 2019). Furthermore, when asymmetric H3 segregation is disrupted, both progenitor germ cell tumors and germ cell loss phenotypes are detected, suggesting that this process is needed for both stem cell maintenance and germ cell differentiation (Xie et al., 2015). The finding of asymmetric histone inheritance in *Drosophila* male GSCs sets a precedent in studying epigenetic inheritance modes in multicellular organisms. The question remained, however, of whether this phenomenon is germ cell-specific or if it serves as a more general mechanism. It also remains unclear whether asymmetric histone inheritance defines distinct cell fates at a single-cell level. Addressing these questions will not only greatly enhance our current understanding of how epigenetic inheritance modes dictate cell fates, but it will also establish new methods that can be used to identify *bona fide* stem cells and/or asymmetrically dividing cells *in vivo*.

To investigate the generality of asymmetric histone inheritance, we used the *Drosophila* intestinal stem cell (ISC) in the midgut as a model system (Micchelli and Perrimon, 2006; Ohlstein and Spradling, 2006). One feature of ISCs is that they can alternate between ACD, which produces a self-renewed ISC and a differentiating enteroblast (EB), and symmetric cell division (SCD), which results in two self-renewed ISCs (Figure 1A) (de Navascues et al., 2012; Martin et al., 2018; O’Brien et al., 2011). As the Notch (N) signaling pathway is critical for cellular differentiation in the ISC lineage, these two modes of cell division can be distinguished at a single-cell resolution using Delta as an ISC-specific marker. We propose that the ISC lineage is a great system to study histone inheritance due to its well-characterized lineage, clearly distinguishable single cell divisions, and abundant stem cells *in vivo*. Using this system, we find that asymmetric histone inheritance applies to this somatic stem cell lineage when ISCs undergo ACD, and that this histone inheritance mode is required for proper cell fate determination as mis-regulation of this process leads to midgut dysplasia and increased ISC-like tumor formation during aging. Collectively, these results demonstrate that asymmetric histone inheritance is a more general feature for asymmetric division of stem cells. This study also offers insight into how histone inheritance influences the establishment of cell identities and how mis-inheritance could lead to diseases such as cancer.

**Figure 1:**
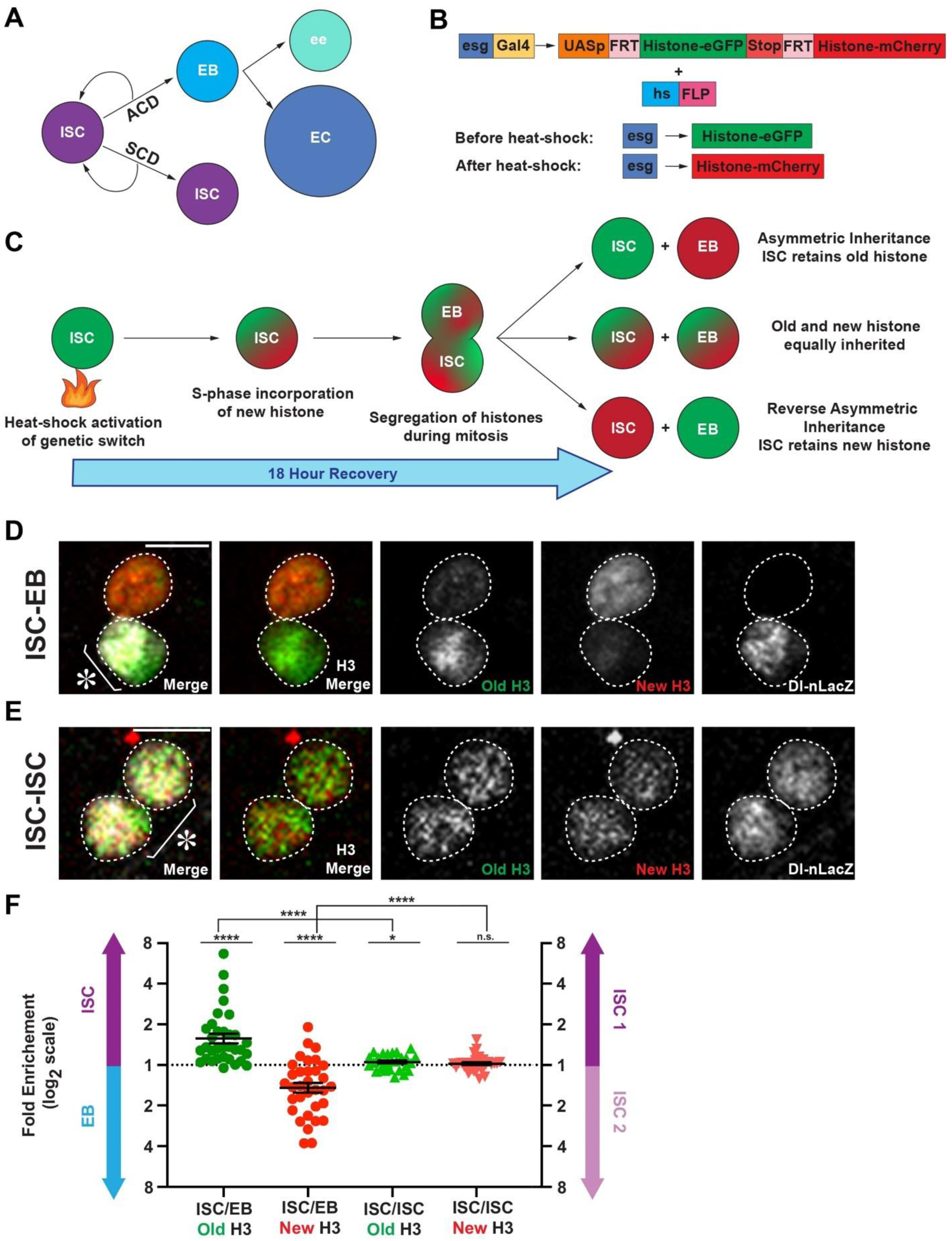
Old and new histone H3 display asymmetric inheritance pattern specifically for asymmetric division of intestinal stem cells. (**A**) A diagram of the ISC lineage. ISCs can asymmetrically divide into a self-renewed ISC and a differentiating enteroblast (EB), or symmetrically divide into two ISCs. EBs can mature into enterocytes (EC) or enteroendocrine cells (ee). (**B**) The dual color labeling system. The histone transgene is driven in the progenitor cells of the ISC lineage using a cell type- and stage-specific *escargot (esg)-Gal4* driver. Before heat shock, the transgene will express eGFP-labeled histones. After heat shock-induced DNA recombination, the transgene will turn off expression of eGFP-labeled histones and turn on expression of mCherry-labeled histones. (**C**) Experimental design. After heat shock, new histones are incorporated genome-wide during S phase, followed by segregation of old and new histones during mitosis. Three possible patterns are shown here: old histones are retained in the ISC (asymmetric), old and new histones are equally inherited (symmetric), or new histones are retained in the ISC (reverse asymmetric). (**D**) Old and new H3 distribution in a post-mitotic ISC-EB pair with Dl-nLacZ labeling to identify the ISC, showing that H3-eGFP (old) is asymmetrically inherited by the ISC, while H3-mCherry (new) is enriched in the EB. Note: all figure panels in this work are maximum intensity projection images. (**E**) Old and new H3 distribution in a post-mitotic ISC-ISC pair with Dl-nLacZ labeling to identify ISCs, showing that both H3-eGFP (old) and H3-mCherry (new) are symmetrically distributed between the two ISCs. (**F**) Quantification of H3-eGFP (old) and H3-mCherry (new) distribution in ISC-EB pairs (avg. log2 ratio for old H3 = 0.65 ± 0.12, avg. log2 ratio for new H3 = −0.56 ± 0.12, n = 33) and ISC-ISC pairs (avg. log2 ratio for old H3 = 0.07 ± 0.03, avg. log2 ratio for new H3 = 0.03 ± 0.03, n = 30). Individual data points and mean values are shown. Error bars represent the standard error of the mean (SEM). **** p < 0.0001, * p <0.05; single sample t test (for normally distributed data) for comparing one dataset to a hypothesized mean of 0 (log2 value representing a 1:1 ratio), or Wilcoxon signed rank test (for skewed data) for comparing one dataset to a hypothesized median of 0 (log2 value representing a 1:1 ratio). Unpaired t test to compare two individual datasets to each other. NS, not significant. Individual data values are shown in Supplemental Table 1. Between the two ISCs, ISC1 has the relatively higher Dl-nLacZ level than ISC2 (Materials and Methods). Scale bar in (**D**) and (**E**): 5 µm; asterisk, ISC side.

## Results

### Old and new H3 display asymmetric inheritance pattern specifically for asymmetric division of intestinal stem cells

To study histone inheritance patterns during ISC divisions, we utilized an optimized dual-color histone labeling and tracking system (Figure 1B), similar to what has been previously used (Tran et al., 2012; Wooten et al., 2019; Xie et al., 2015). Here, the expression of labeled histones was driven by the cell type-specific *escargot-Gal4* (*esg-Gal4*) driver, which turns on the *UAS-histone* transgene exclusively in ISCs and EBs in this adult stem cell lineage (Micchelli and Perrimon, 2006). After a heat-shock induced switch from eGFP to mCherry labeled histone expression, ISCs were allowed to undergo a complete round of DNA replication after a prolonged recovery time for 18 hours, visualized by robust incorporation of new canonical histones (Figure 1C). The co-labeled ISCs with both eGFP (old histones) and mCherry (new histones) signals then entered the subsequent mitosis, where sister chromatids and the incorporated histones are segregated to the daughter cells.

We first examined eGFP-labeled old histone *versus* mCherry-labeled new histone inheritance patterns in the two daughter cells that result from a recent ISC division (Figure 1C). As ISCs are the only mitotically active cells that express this histone transgene (Jiang et al., 2009; Micchelli and Perrimon, 2006; Ohlstein and Spradling, 2006), post-mitotic pairs can be precisely identified as two neighboring cells both carrying new histones, which should result from the previous S-phase-dependent new histone incorporation and the subsequent ISC division. Two potential caveats are that first, ISCs could enter the subsequent S-phase after one cell division and incorporate more new histones; and second, that EBs could begin to mature and endocycle, incorporating more new histones (Micchelli and Perrimon, 2006; Ohlstein and Spradling, 2006, 2007). To exclude these situations, a 30-minute pulse of the nucleoside analog EdU (5-ethynyl-2’-deoxyuridine) was used to identify ISCs undergoing the next S-phase and EBs that are endocycling (Materials and Methods). Additionally, a Delta-nuclear lacZ (Dl-nLacZ) reporter was used to distinguish ACD-derived ISC-EB pairs from SCD-derived ISC-ISC pairs, as this reporter is specifically expressed in ISCs (Beebe et al., 2010; Jiang et al., 2009; Zeng et al., 2010).

Using these methods to precisely pinpoint the daughter cells from different cell division modes and at the proper cell cycle phase, we found that old and new H3 displayed an asymmetric pattern in the ISC-EB pair (Figure 1D). In contrast, H3 was inherited in a more symmetric manner in ISC-ISC pairs (Figure 1E). Quantification of these results showed that in ISC-EB pairs, old H3 was preferentially inherited by the ISC while new H3 was enriched in the EB. In contrast, both old and new H3 were distributed almost equally between the two daughter ISCs (Figure 1F). Similar results were obtained using the same histone labeling strategy in a parallel experimental regime, but with an antibody that recognizes the Delta protein (Supplementary Figure 1A-C) (Bardin et al., 2010; Obniski et al., 2018; Ohlstein and Spradling, 2007). Thus, the asymmetric histone inheritance pattern observed is specific to ISC-EB pairs resulting from asymmetric ISC divisions, as ISC-ISC pairs resulting from symmetric ISC divisions inherited both old and new histones more equally. This cellular specificity demonstrates that asymmetric histone inheritance is specific to asymmetrically dividing stem cells, where distinct cell fates are generated through one cell division.

### Old and new H4 display asymmetric inheritance pattern while old and new H2A display symmetric inheritance pattern for asymmetric division of intestinal stem cells

In eukaryotic cells, every ∼147 bp of double helix DNA wraps around an octamer structure composed of histone H2A, H2B, H3, and H4, each in two copies. During DNA replication, preexisting histone octamers must be disassembled and reincorporated into the duplicated sister chromatids [reviewed in (Serra-Cardona and Zhang, 2018; Snedeker et al., 2017; Stewart-Morgan et al., 2020; Xu and Zhu, 2010)]. It has been demonstrated that preexisting H3 and H4 are reincorporated as a tetramer, while old H2A and H2B are reincorporated as two dimers following their dissociation from DNA strands during replication (Jackson, 1988; Jackson and Chalkley, 1981; Katan-Khaykovich and Struhl, 2011; Russev and Hancock, 1981; Xu et al., 2010). To investigate the molecular specificity of asymmetric histone inheritance, we further examined the inheritance modes of histones H4 and H2A using similar dual-color labeling strategies (Figure 1B). We found that old and new H4 displayed an asymmetric inheritance pattern in post-mitotic ISC-EB pairs derived from ACD of ISCs, shown by the asymmetric distribution of Dl-nLacZ reporter as described above (Figure 2A). Similar to H3, the asymmetric inheritance pattern of old *versus* new H4 was specific to the ACD mode of ISCs, as the SCD mode that leads to ISC-ISC pairs with comparable Dl-nLacZ levels showed a more symmetric distribution of old *versus* new H4 (Figure 2B). Consistent with the H3 results, quantification of these results showed that old H4 was preferentially inherited by the self-renewed ISC while new H4 was enriched in the differentiating EB. In contrast, both old and new H4 were distributed almost equally between the two daughter ISCs (Figure 2C).

**Figure 2:**
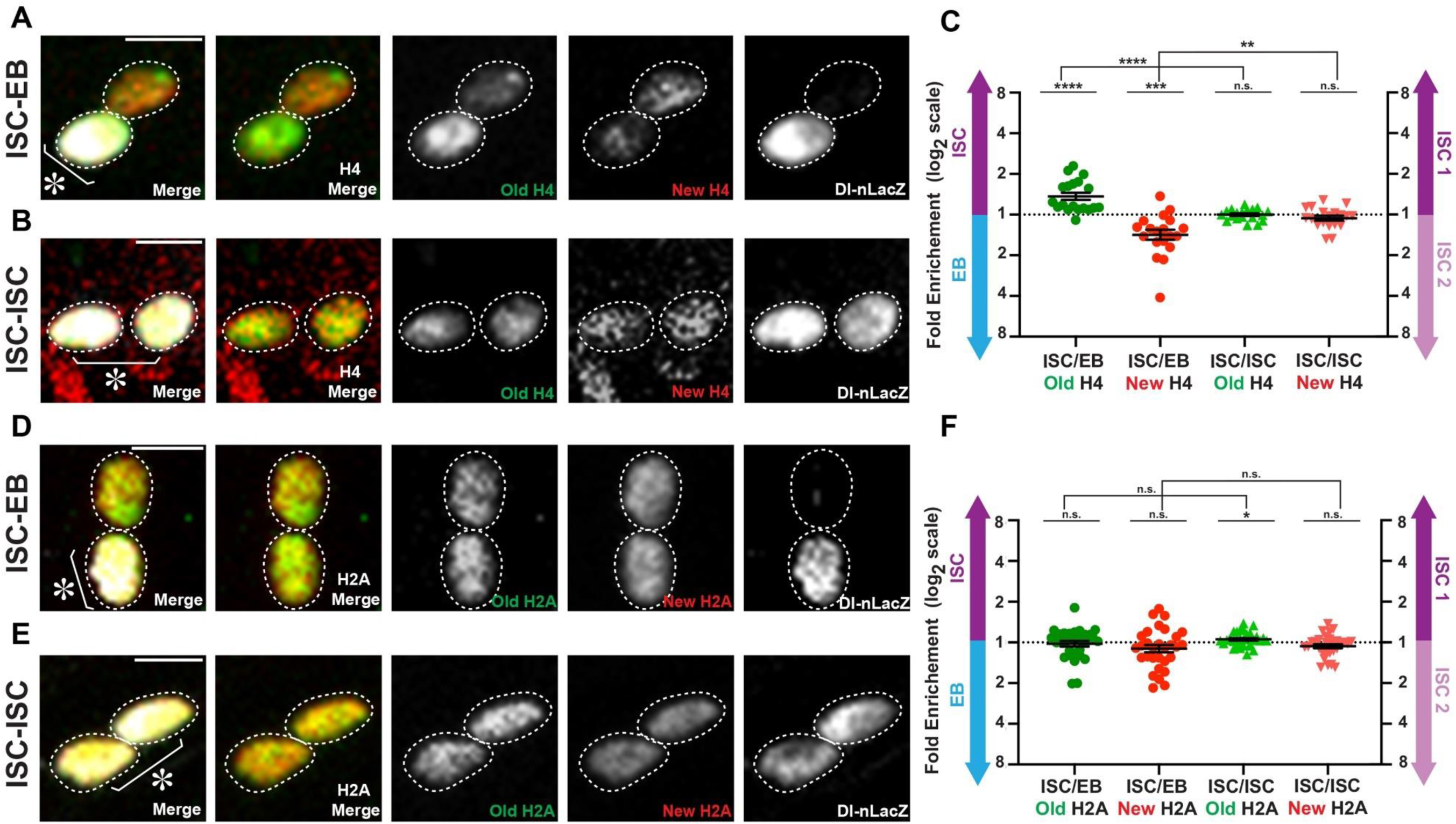
Old and new H4 display asymmetric inheritance pattern while old and new H2A display symmetric inheritance pattern for asymmetric division of intestinal stem cells. (**A**) Old and new H4 distribution in a post-mitotic ISC-EB pair with Dl-nLacZ labeling to identify the ISC, showing that H4-eGFP (old) is asymmetrically inherited by the ISC, while H4-mCherry (new) is enriched toward the EB. (**B**) Old and new H4 distribution in a post-mitotic ISC-ISC pair with Dl-nLacZ labeling to identify ISCs, showing that both H4-eGFP (old) and H4-mCherry (new) are symmetrically distributed between the two ISCs. (**C**) Quantification of H4-eGFP (old) and H4-mCherry (new) distribution in ISC-EB pairs (avg. log2 ratio for old H4 = 0.45 ± 0.09, avg. log2 ratio for new H4 = −0.50 ± 0.12, n = 19) and ISC-ISC pairs (avg. log2 ratio for old H4 = 0.00 ± 0.03, avg. log2 ratio for new H4 = −0.09 ± 0.06, n = 19). Individual data points and mean values are shown. Error bars represent SEM. **** p < 0.0001, *** p < 0.001, ** p < 0.01; single sample t test (for normally distributed data) for comparing one dataset to a hypothesized mean of 0 (log2 value representing a 1:1 ratio), or Wilcoxon signed rank test (for skewed data) for comparing one dataset to a hypothesized median of 0 (log2 value representing a 1:1 ratio). Unpaired t test to compare two individual datasets to each other. NS, not significant. Individual data values are shown in Supplemental Table 1. (**D**) Old and new H2A distribution in a post-mitotic ISC-EB pair with Dl-nLacZ labeling to identify the ISC, showing that H2A-eGFP (old) and H2A-mCherry (new) are symmetrically inherited in the ISC and EB. (**E**) Old and new H2A distribution in a post-mitotic ISC-ISC pair with Dl-nLacZ labeling to identify ISCs, showing that both H2A-eGFP (old) and H2A-mCherry (new) are symmetrically distributed between the two ISCs. (**F**) Quantification of H2A-eGFP (old) and H2A-mCherry (new) distribution in ISC-EB pairs (avg. log2 ratio for old H2A = −0.03 ± 0.07, avg. log2 ratio for new H2A = −0.15 ± 0.09, n = 30) and ISC-ISC pairs (avg. log2 ratio for old H2A = 0.08 ± 0.03, avg. log2 ratio for new H3 = – 0.09 ± 0.05, n = 30). Individual data points and mean values are shown. Error bars represent SEM. * p <0.05; single sample t test (for normally distributed data) for comparing one dataset to a hypothesized mean of 0 (log2 value representing a 1:1 ratio), or Wilcoxon signed rank test (for skewed data) for comparing one dataset to a hypothesized median of 0 (log2 value representing a 1:1 ratio). Unpaired t test to compare two individual datasets to each other. NS, not significant. Individual data values are shown in Supplemental Table 1. Scale bar in (**A**), (**B**), (**D**), and (**E**): 5 µm; asterisk, ISC side.

Notably, the ratios reflecting the degree of asymmetry for old and new H4 (∼1.37-fold for old H4 and ∼1.41-fold for new H4) is less than that of old and new H3 (∼1.57-fold for old H3 and ∼1.47-fold for new H3) in the ISC-EB pairs derived from ISC ACD (Figure 2C vs. Figure 1F), despite all of these ratios being significantly different from a symmetrical ratio (1:1) and the ratios from ISC-ISC pairs resulted from ISC SCD (see *P* values in Figure 2C). This difference could be due to the fact that in higher eukaryotes including *Drosophila*, H4 partners with H3 and the histone variants, such as H3.3 and the centromere-specific CENP-A. Both H3.3 and CENP-A are not incorporated restrictively in a replication-dependent manner [reviewed in (Szenker et al., 2011)]. The incorporation of H3.3 is more dependent on transcription (Ahmad and Henikoff, 2002a, b; Tagami et al., 2004). Even during replication, old H3-H4 remain as tetramers but the H3.3-H4 tetramers likely split into two dimers (Xu et al., 2010), which would affect their re-incorporation and the overall inheritance pattern. For CENP-A, it has been shown that old CENP-A is preferentially retained by ISC in ACD (Garcia Del Arco et al., 2018). The transcription-dependent H3.3-H4 deposition, the possibly more symmetric re-incorporation of H3.3-H4 during replication, and the incorporation timing and mode of CENP-A-H4 together may contribute to this slight difference of the ratios between H3 and H4.

Next, we examined the inheritance mode of old *versus* new H2A as a representative for the canonical histone dimer. Different from H3 and H4, old and new H2A showed a symmetric inheritance pattern in all post-mitotic daughter cells of ISCs, regardless of whether they were an asymmetric ISC-EB pair (Figure 2D, 2F) or a symmetric ISC-ISC pair (Figure 2E, 2F). Quantification for old and new H3A showed no significant difference from a symmetrical pattern (a 1:1 ratio) for ACD of ISCs, and there was no significant difference between ACD and SCD of ISCs (see *P* values in Figure 2F). Similar results were obtained using the same histone labeling strategy in a similar experimental regime, using an antibody that recognizes the Delta protein (Supplementary Figure 1D-F). This molecular difference between H2A and H3/H4 is consistent with previous biochemical results and recent discoveries in the *Drosophila* male GSCs (Tran et al., 2012; Wooten et al., 2019). This molecular specificity of asymmetric H3/H4 inheritance is likely because re-incorporation of the old (H3-H4)2 tetramers greatly enhances the asymmetric distribution of old *versus* new H3/H4 between sister chromatids during DNA replication.

### Asymmetric old and new H3 segregation and distribution during mitosis of intestinal stem cells

Through post-mitotic daughter cell pairs, we found that old *versus* new H3 and H4 are asymmetrically distributed in ISC-EB pairs and symmetrically distributed in ISC-ISC pairs, while old *versus* new H2A are distributed symmetrically in both ISC-EB and ISC-ISC pairs. To understand these patterns in relation to cell cycle progression, we studied old *versus* new histone distribution at different mitotic stages of ISCs. We reason that in order to achieve the asymmetric inheritance pattern in ISC-EB pairs after mitosis, the differential distribution between old and new histones should have been established prior to mitosis. Consequently, their differential localization could be detectable as the chromosomes condense in prophase to prometaphase and as sister chromatids segregate in anaphase to telophase.

As ISCs are the only mitotically active cells in the entire lineage (Jiang et al., 2009; Micchelli and Perrimon, 2006; Ohlstein and Spradling, 2006), they can be readily identified using a mitotically enriched H3S10ph mark (phosphorylation at Serine 10 of H3) (Hendzel et al., 1997). When old and new H3 distribution patterns were examined in anaphase and telophase ISCs, we visualized asymmetric segregation patterns (Figure 3A). In contrast, old and new H2A displayed a more symmetric segregation pattern in anaphase and telophase ISCs (Figure 3B). We then quantified the ratios of the old histone between the two sets of sister chromatids at anaphase and telophase (Figure 3C) (Xie et al., 2015). Based on this quantification, old H3 tended to be enriched towards one set of sister chromatids, whereas old H2A distributed more equally between the two sets of sister chromatids. Using different thresholds to classify the degree of old H3 asymmetry, approximately 40% of mitotic ISCs showed a high degree of asymmetry (>1.4-fold difference), ∼40% showed a medium level of asymmetry (1.2-1.4-fold difference), and ∼20% demonstrated a symmetric pattern [<1.2-fold difference, Figure 3D and (Ranjan et al., 2019)]. Notably, these ratios are consistent with other published results measuring the percentages of ACD *versus* SCD of ISCs using different cellular markers or criteria (de Navascues et al., 2012; Goulas et al., 2012; Jin et al., 2017; O’Brien et al., 2011; Tian and Jiang, 2014). In contrast, 95% of all quantified mitotic ISCs showed a symmetric segregation pattern of old H2A in anaphase or telophase ISCs, further confirming the molecular specificity of this asymmetric histone inheritance pattern.

**Figure 3:**
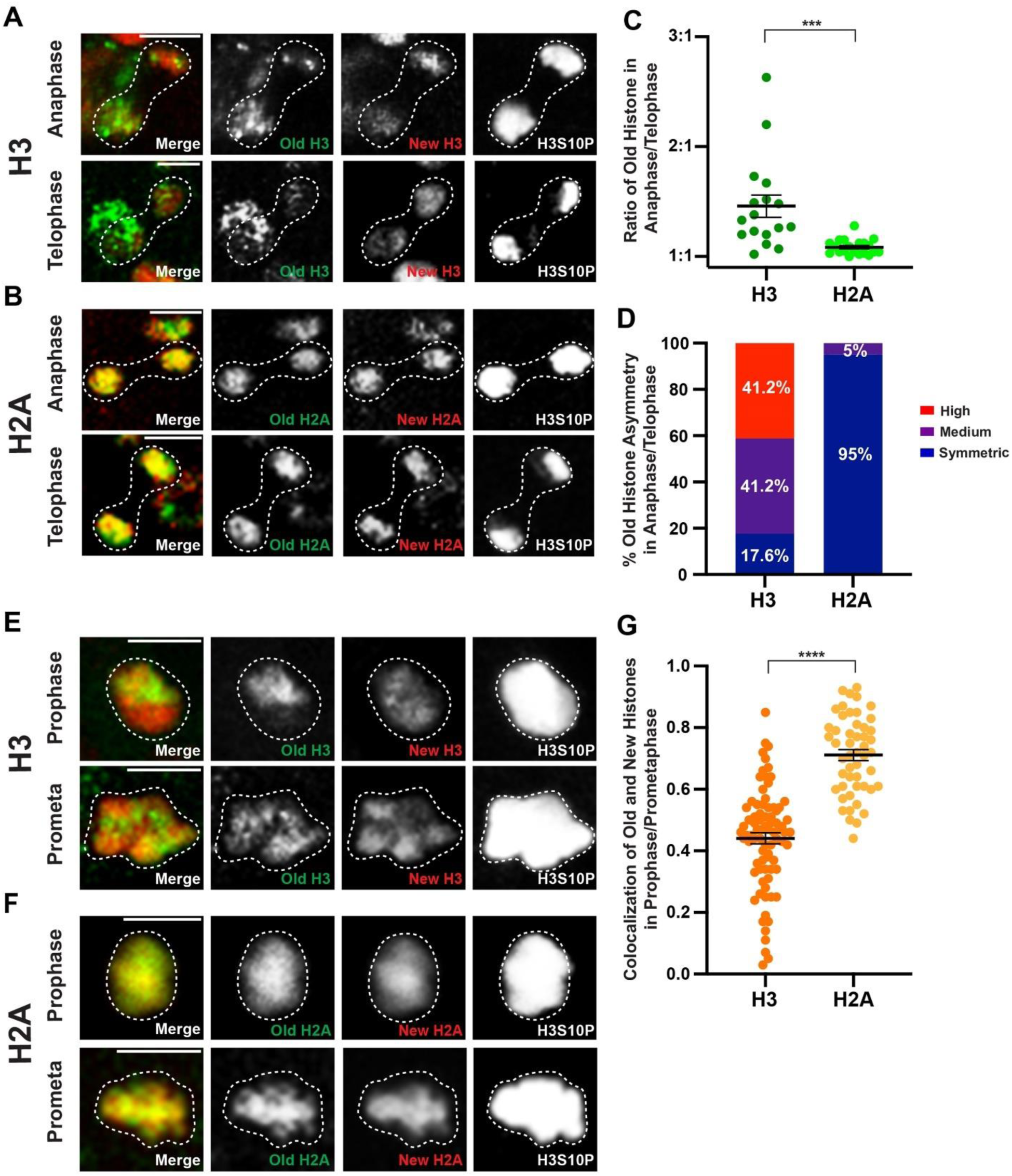
Asymmetric old and new H3 distribution and segregation during mitosis of intestinal stem cells. (**A**) Anaphase and telophase ISCs showing asymmetric segregation of H3-eGFP (old) and H3-mCherry (new) between the two sets of sister chromatids at the opposite poles of the ISC. Note: For the telophase image, sister chromatids from the ISC are outlined, the H3-eGFP signal outside the outline is from a neighboring cell shown up in this maximum intensity projection image. (**B**) Anaphase and telophase ISCs showing symmetric segregation of H2A-eGFP (old) and H2A-mCherry (new) between the two sets of sister chromatids at the opposite poles of the ISC. (**C**) Quantification of old H3 distribution between the two sets of sister chromatids in anaphase and telophase ISCs (avg. old H3 = 1.46 ± 0.10, n = 17) or H2A (avg. old H2A = 1.08 ± 0.02, n = 20). Old histone signals are used here, due to potential complications with new histone signals that have been discussed in this work and previously (Xie et al., 2015). Individual data points and mean values are shown. Error bars represent SEM. *** p < 0.001; unpaired t test to compare two individual datasets to each other. Individual data values are shown in Supplemental Table 2. (**D**) Percentage of different categories of old histone asymmetry for H3 (n = 17) and H2A (n = 20), where the three categories are symmetric (<1.2-fold), medium asymmetric (1.2- to 1.4-fold), and highly asymmetric (>1.4-fold). (**E**) Old and new H3 distribution in prophase and prometaphase ISCs, showing separable domains of H3-eGFP (old) and H3-mCherry (new). (**F**) Old and new H2A distribution in prophase and prometaphase ISCs, showing largely overlapping signals between H2A-eGFP (old) and H2A-mCherry (new). (**G**) Quantification of colocalization between eGFP- and mCherry-tagged H3, as well as between eGFP- and mCherry-tagged H2A in prophase and prometaphase ISCs. Pearson’s correlation coefficients are recorded, where 1 represents complete colocalization and 0 stands for no colocalization. eGFP- and mCherry-tagged H3 showed significantly less colocalization (avg. Pearson’s correlation coefficient for H3 = 0.44 ± 0.02, n = 81) when compared to eGFP- and mCherry-tagged H2A (avg. Pearson’s correlation coefficient for H2A = 0.71 ± 0.02, n = 50). Individual data points and mean values are shown. Error bars represent SEM. **** p < 0.0001; unpaired t test to compare two individual datasets to each other. Individual data values are shown in Supplemental Table 3. Scale bar in (**A**), (**B**), (**E**): 5 µm.

Interestingly, separable domains of old and new H3 could already be visualized during prophase and prometaphase, when chromosomes undergo condensation (Figure 3E). However, this separation was not as evident for old and new H2A, which displayed a more overlapping pattern in prophase and prometaphase ISCs (Figure 3F). We used Pearson’s correlation analysis to determine the colocalization between old and new histones for both H3 and H2A, where a correlation coefficient of 1 indicates perfect colocalization and 0 indicates no colocalization. The average correlation coefficient for old and new H3 was 0.44, while the average coefficient for old and new H2A was 0.71, which are significantly different (Figure 3G). Similar results were obtained by using the Spearman correlation coefficient to determine colocalization of old *versus* new H3 and H2A, respectively (Supplementary Figure 2). Taken together, these results suggest that old and new H3 are differentially incorporated onto sister chromatids prior to mitosis, while old and new H2A are more uniformly incorporated.

### Expression of a mutant histone H3T3A disrupts histone inheritance patterns

We have previously identified a mitosis-specific phosphorylation at Threonine 3 of H3 (H3T3ph) that can distinguish sister chromatids enriched with old *versus* new H3, consistent with biochemistry results (Lin et al., 2016). Differential H3T3ph at old H3-*versus* new H3-enriched sister chromatids coordinate their proper recognition and segregation during ACD of male GSCs. By mutating the T3 residue on the H3 tail to an unphosphorylatable Alanine (H3T3A), the asymmetric segregation of old and new H3 become randomized. This mutation also resulted in cellular defects including early germline tumors and germ cell loss, indicating that the mis-regulation of asymmetric histone inheritance might affect proper cell identity establishment through ACD of GSCs (Xie et al., 2015). To further study this histone mutant and its potential effects on both histone inheritance and distinct cell fate establishment in the ISC lineage, we expressed H3T3A using the same *esg-Gal4* driver, as described previously.

We first detected a significant increase in the percentage of ISC-ISC pairs in the *esg*>H3T3A mutant histone-expressing midgut as compared to the *esg*>H3 expressing midgut (Figure 4A-C). Here, both fly lines were cultured under the same conditions. These results suggest that expressing the H3T3A mutant histone results in a higher occurrence of symmetric ISC divisions than expressing wild-type H3. When the dual color switching experiments using the H3T3A-containing transgene were performed (Figure 1B), both ISC-ISC and ISC-EB pairs using the Dl-nLacZ reporter could be identified (Figure 4B-C). However, in *esg*>H3T3A midgut a lower percentage of ISC-EB pairs was detected compared to *esg*>H3 midgut (Figure 4A). In these ISC-EB pairs, the Dl-nLacZ-enriched ISC cell still displayed a higher level of old H3T3A than the Dl-nLacZ-deprived EB cell, whereas the EB cell was more enriched with new H3T3A (Figure 4C). These observations suggest that old *versus* new H3T3A are still differentially deposited onto sister chromatids prior to mitosis and could be asymmetrically inherited, albeit less frequently.

**Figure 4:**
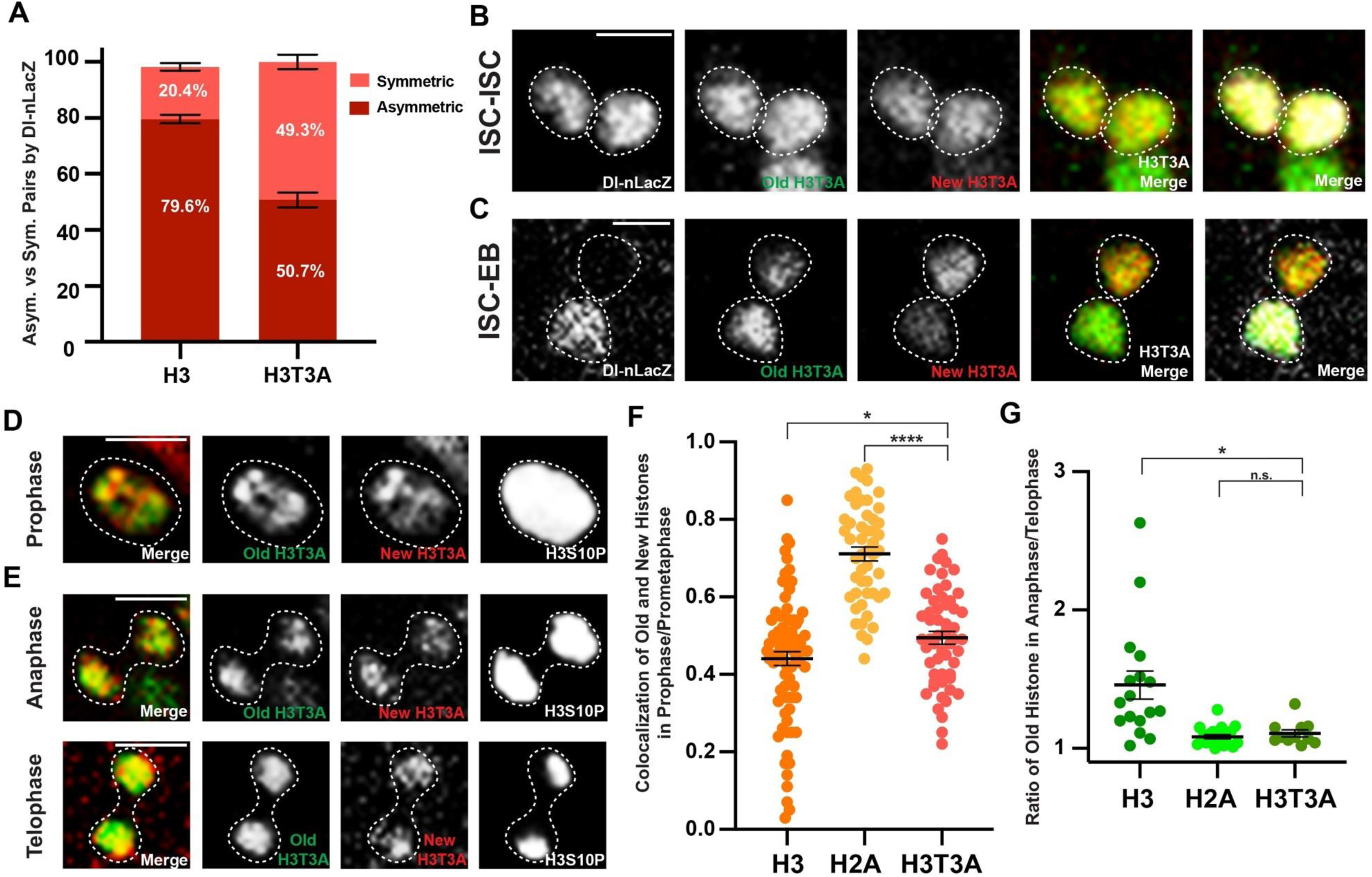
Expression of a mutant histone H3T3A disrupts histone inheritance patterns and increases symmetric ISC divisions. (**A**) Quantification of asymmetric (ISC-EB) and symmetric (ISC-ISC) pairs in wild-type H3-expressing midguts (avg. asymmetric = 79.6% ± 1.5%, avg. symmetric = 20.4% ± 1.4% N = 6 midguts, n = 2341 pairs) and H3T3A-expressing midguts (avg. asymmetric = 50.7% ± 2.6%, avg. symmetric = 49.3% ± 2.6%, N = 6 midguts, n = 2747 pairs). H3T3A midguts have a significant increase in symmetric ISC-ISC pairs (two-sample t test, p < 0.0001). Individual data values are shown in Supplemental Table 4. (**B**) In a post-mitotic ISC-ISC pair labeled with comparable Dl-nLacZ, both H3T3A-GFP (old) and H3T3A-mKO (new) are symmetrically distributed between the two ISCs. (**C**) In a post-mitotic ISC-EB pair with ISC labeled with higher Dl-nLacZ, H3T3A-GFP (old) is enriched in the ISC, while H3T3A-mKO (new) is more toward the EB. (**D**) Old and new H3T3A distribution in prophase and prometaphase ISCs, showing separable domains between H3T3A-GFP (old) and H3T3A-mKO (new). (**E**) Anaphase and telophase ISCs showing symmetric segregation of H3T3A-GFP (old) and H3T3A-mKO (new) between the two sets of sister chromatids at the opposite poles of the ISC. (**F**) Quantification of colocalization between H3T3A-GFP (old) and H3T3A-mKO (new) in prophase and prometaphase ISCs using Pearson’s correlation coefficients. The H3 and H2A data are from Figure 3G for direct comparison. Old and new H3T3A distribution is similar to that of H3 (avg. Pearson’s correlation coefficient for H3T3A = 0.49 ± 0.02, n = 55). Individual data points and mean values are shown. Error bars represent SEM. **** p < 0.0001, * p < 0.05; unpaired t test to compare two individual datasets to each other. Individual data values are shown in Supplemental Table 3. (**G**) Quantification of old histone distribution between the two sets of sister chromatids for anaphase and telophase ISCs expressing H3T3A (avg. old H3T3A = 1.11 ± 0.03, n = 11). The H3 and H2A data are from Figure 3C for direct comparison. Old and new H3T3A segregation pattern is similar to H2A, showing more symmetric pattern. Individual data points and mean values are shown. Error bars represent SEM. * p < 0.05; unpaired t test to compare two individual datasets to each other. NS, not significant. Individual data values are shown in Supplemental Table 2. Scale bar in (**B**), (**C**), (**D**), (**E**): 5 µm; asterisk, ISC side.

To further investigate this phenomenon, we studied the distribution of old *versus* new H3T3A histones throughout mitosis in ISCs. When examining prophase and prometaphase ISCs, distinct old *versus* new H3T3A domains could still be detected, similar to the pattern found in old *versus* new H3 at the same phases in mitotic ISCs (Figure 4D vs. Figure 3E). When conducting Pearson’s correlation analyses, the average correlation coefficient of 0.49 for old *versus* new H3T3A resembled the average correlation coefficient of 0.44 for old *versus* new H3, both of which were significantly lower than the average correlation coefficient of 0.71 for old *versus* new H2A (Figure 4F). The Spearman correlation coefficient yielded similar results from assessing the colocalization of old *versus* new H3T3A (Supplementary Figure 2). These results indicate that old and new H3T3A are still differentially incorporated onto sister chromatids prior to mitosis, just like wild-type H3.

However, when compared to wild-type old H3, old H3T3A was distributed more equally between the two segregated sets of sister chromatids in anaphase and telophase in ISCs (Figure 4E vs. Figure 3A). When quantified, the segregation pattern of old H3T3A was more similar to that of H2A than to H3 (Figure 4G). Collectively, these data suggest that old *versus* new H3T3A are differentially incorporated into sister chromatids prior to mitosis, similar to H3; however, this mutation disrupts the differential recognition of sister chromatids in mitotic ISCs, leading to symmetric segregation of old and new H3T3A at anaphase and telophase. Notably, these results are consistently in accordance with the previous findings regarding the incorporation and segregation modes of old *versus* new H3T3A in the male GSCs (Xie et al., 2015), suggesting a conserved underlying mechanism in both stem cell systems.

### Mutant histone H3T3A leads to ISC overproliferation and increased intestinal tumors

In the H3T3A-expressing ISC lineage, the occurrence of ISC-ISC pairs increased to 49.3%, significantly higher than 20.4% in the H3-expressing ISC lineage (Figure 4A). Examining the morphology of the H3T3A-expressing midgut revealed different distribution of ISCs that express the Delta-nLacZ reporter (Figure 5A, 5B): In the wild-type H3-expressing midgut, Dl-nLacZ positive cells could be detected as single-cell colonies in an evenly dispersed manner. Two Dl-nLacZ positive cells were only adjacent to each other occasionally as a result of an ISC symmetric division (Figure 5A). This is consistent with previous reports of the ISC distribution pattern in the midgut (Ahmed et al., 2020; Ohlstein and Spradling, 2007). In contrast, a disorganized pattern of Dl-nLacZ positive cells could be detected in the H3T3A-expressing midgut, with an appearance of increased ISC clusters with more than one cell (Figure 5B). Using a clustering analysis, we quantified the relative frequencies of cell cluster(s) with 1-, 2-, or ≥3 Dl-nLacZ positive cells (Figure 5C). This analysis revealed a substantial increase of 2- and ≥3 Dl-nLacZ cell clusters in the H3T3A-expressing midgut as compared to the H3- expressing midgut, suggesting an increased ratio of SCD and potentially an overproliferation of ISC-like cells in the H3T3A-expressing midgut (Figure 5D). This increase is at the expense of single Dl-nLacZ positive cells, which displayed a significant decrease from 77.9% in H3- expressing midguts to 50.4% in H3T3A-expressing midguts (Figure 5E). These results further support increased SCD events detected in the H3T3A-expressing midgut (Figure 4A).

**Figure 5:**
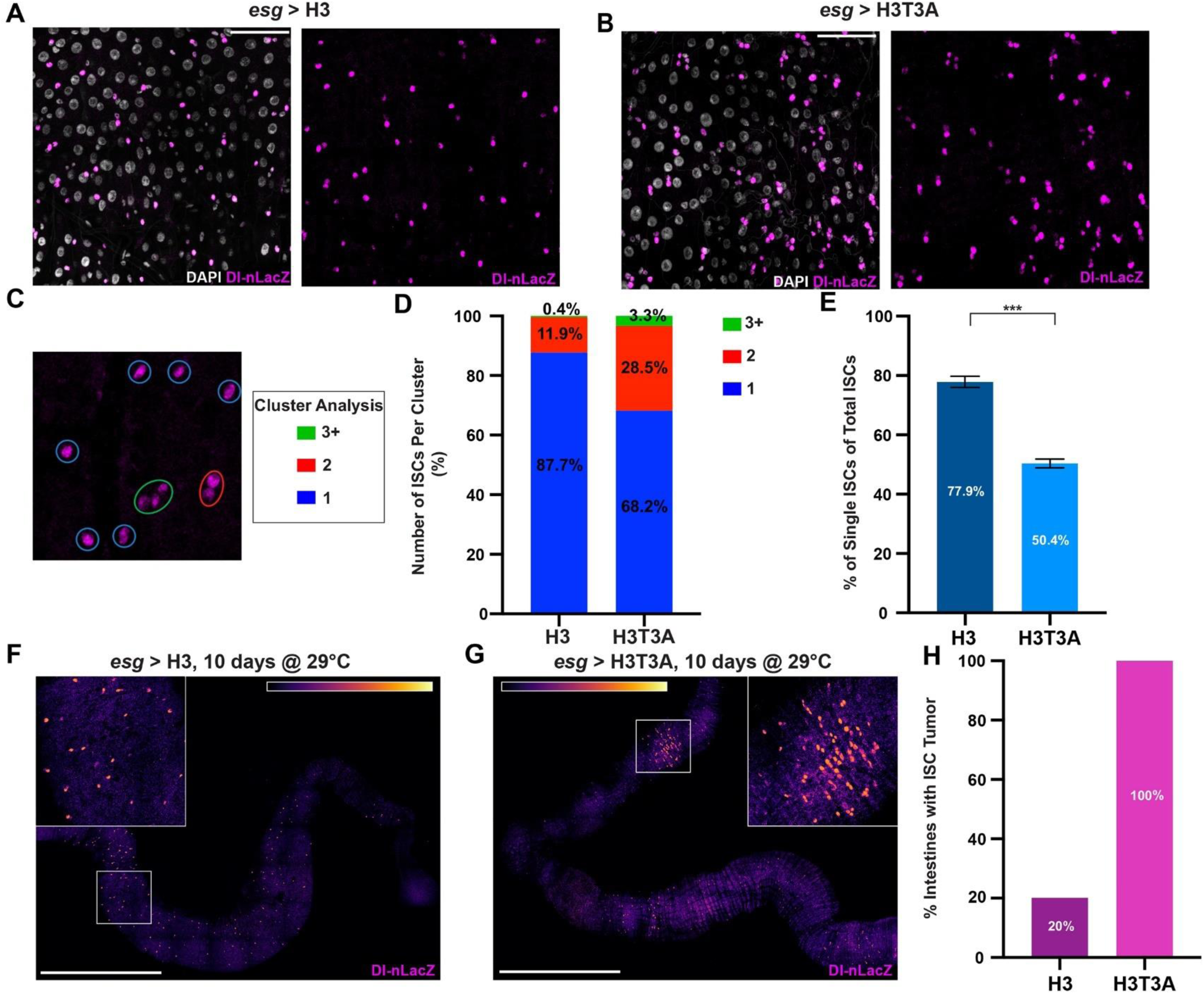
Mutant histone H3T3A leads to ISC overproliferation and increased intestinal tumors. (**A**) Distribution of the Dl-nLacZ-positive (magenta) ISCs in an H3-expressing midgut. ISCs are well spaced and interspersed within the intestinal epithelium. (**B**) Distribution of the Dl-nLacZ-positive (magenta) ISCs in an H3T3A-expressing midgut. ISCs are unevenly distributed in clusters with two or more ISCs. DAPI: white in (**A-B**). (**C**) Cluster analyses show representative 1-, 2-, or ≥3-ISC clusters. (**D**) Percentages of clusters with 1-, 2-, or ≥3-ISC clusters for H3-expressing midguts (avg. percentage of 1-ISC= 87.7% ± 1.1%, avg. percentage of 2-ISC= 11.9% ± 0.9%, avg. percentage of ≥3-ISC= 0.4% ± 0.2%, N = 4 midguts, n = 1482 clusters) and H3T3A-expressing midguts (avg. percentage of 1-ISC= 68.2% ± 1.2%, avg. percentage of 2-ISC= 28.5% ± 0.9%, avg. percentage of ≥3-ISC= 3.3% ± 0.4%, N = 5 midguts, n = 1862 clusters). Comparing H3-expressing midguts with H3T3A-expressing midguts, there was a significant decrease in 1-ISC clusters (p < 0.0001), and a significant increase in 2-ISC clusters (p < 0.0001) and 3-ISC clusters (p < 0.0001), all by two sample t test. Individual data values are shown in Supplemental Table 5. (**E**) Percentages of single ISCs out of total ISCs for H3-expressing midguts (avg. single ISCs = 77.9% + 1.9%, N = 4 midguts, n = 1674 ISCs) and H3T3A-expressing midguts (avg. single ISCs = 50.4% ± 1.5%, N = 5 midguts, n = 2522 ISCs). There are fewer single ISCs in H3T3A-expressing midguts compared to H3-expressing midguts (two sample t test, *** p < 0.001) (**F**) H3-expressing midgut aged for 10 days at 29°C. Midgut looks largely normal, with an even distribution of mostly single ISCs throughout the tissue. (**G**) H3T3A-expressing midgut aged for 10 days at 29°C. ISCs are unevenly disbursed throughout the midgut, and a large region with overproliferated ISCs can be visualized, classified as an intestinal tumor. (**H**) Percentage of H3-expressing midguts (n = 10) and H3T3A-expressing midguts (n = 12) with tumor(s). Scale bars in (**A**) and (**B**): 50 µm, in (**F**) and (**G**): 500 µm.

Previously it has been shown that aging has a profound effect on ISC behavior, especially the tendency to develop midgut tumors (Biteau et al., 2008; Choi et al., 2008; Regan et al., 2016). Next, we studied the effects of aging on H3T3A-expressing midgut to test whether H3T3A could enhance tumor growth during aging. When *esg>*H3 and *esg>*H3T3A flies were cultured under the same condition for ten days at 29°C to accelerate the aging process (Cohet, 1975; Miquel et al., 1976), a high density of Dl-nLacZ positive cells appeared in the H3T3A-expressing midguts, resulting in ISC-like tumors. In contrast, the control H3-expressing midguts mostly maintained a normal distribution pattern of Dl-nLacZ positive ISCs, shown as single colonies in a well-spaced manner (Figure 5F, 5G). When quantified, every single H3T3A-expressing midgut developed ISC-like tumors, indicating a full penetrance of this phenotype. In contrast, only 20% of H3-expressing midguts showed such a phenotype (Figure 5H). In summary, these results from the ISC lineage demonstrate that the H3T3A mutation causes mis-regulation of histone inheritance and failure in proper cell fate determination, pointing to the notion that differential histone inheritance defines distinct daughter cell identities arising from ACD of ISCs. Notably, the consistency of these results with previous findings regarding H3T3A in the male GSC (Xie et al., 2015) suggests potential conserved mechanisms underlying asymmetric histone inheritance in these two adult stem cell systems.

## Discussion

A long-standing critical biological question is how distinct cell fates are established, maintained, and changed by epigenetic mechanisms at the single-cell level in multicellular organisms. Here, using the *Drosophila* ISC lineage as a model system, we studied the inheritance of different canonical histones H3, H4, and H2A during both ACD and SCD of ISCs. We found that histone inheritance is asymmetric during ACD, where the self-renewed ISC inherited more pre-existing (old) histones, and the differentiating daughter cell inherited more newly synthesized (new) histones. In contrast, histones were inherited symmetrically between the two identical daughter ISCs resulting from SCD of ISC (Figure 6A). These findings directly link the asymmetric histone inheritance mode with the establishment of distinct cell identities after one cell division. Furthermore, we found that this asymmetric inheritance mode is specific to H3 and H4 histones (Figure 6A), as old and new H2A histones are symmetrically inherited even during ACD of ISCs (Figure 6B). This molecular specificity could be explained by the incorporation of old H3 and H4 into chromatin as a (H3-H4)2 tetramer, while old H2A and H2B are incorporated as (H2A-H2B) dimers (Jackson, 1988; Jackson and Chalkley, 1981; Katan-Khaykovich and Struhl, 2011; Russev and Hancock, 1981; Xu et al., 2010). Since H3 and H4 carry most of the post-translational modifications that influence gene expression (Allis and Jenuwein, 2016; Huang et al., 2014; Kouzarides, 2007; Young et al., 2010), the asymmetric inheritance of old and new (H3-H4)2 serves as an elegant mechanism for establishing distinct epigenomes while still maintaining the same genetic information in the daughter cells that arise from stem cell ACD, leading to differential gene expression programs and distinct cell fates. Together, these results indicate that the histone inheritance mode is directly linked with cell fate: asymmetric mode with distinct cell fates while symmetric mode with identical cell fate.

**Figure 6:**
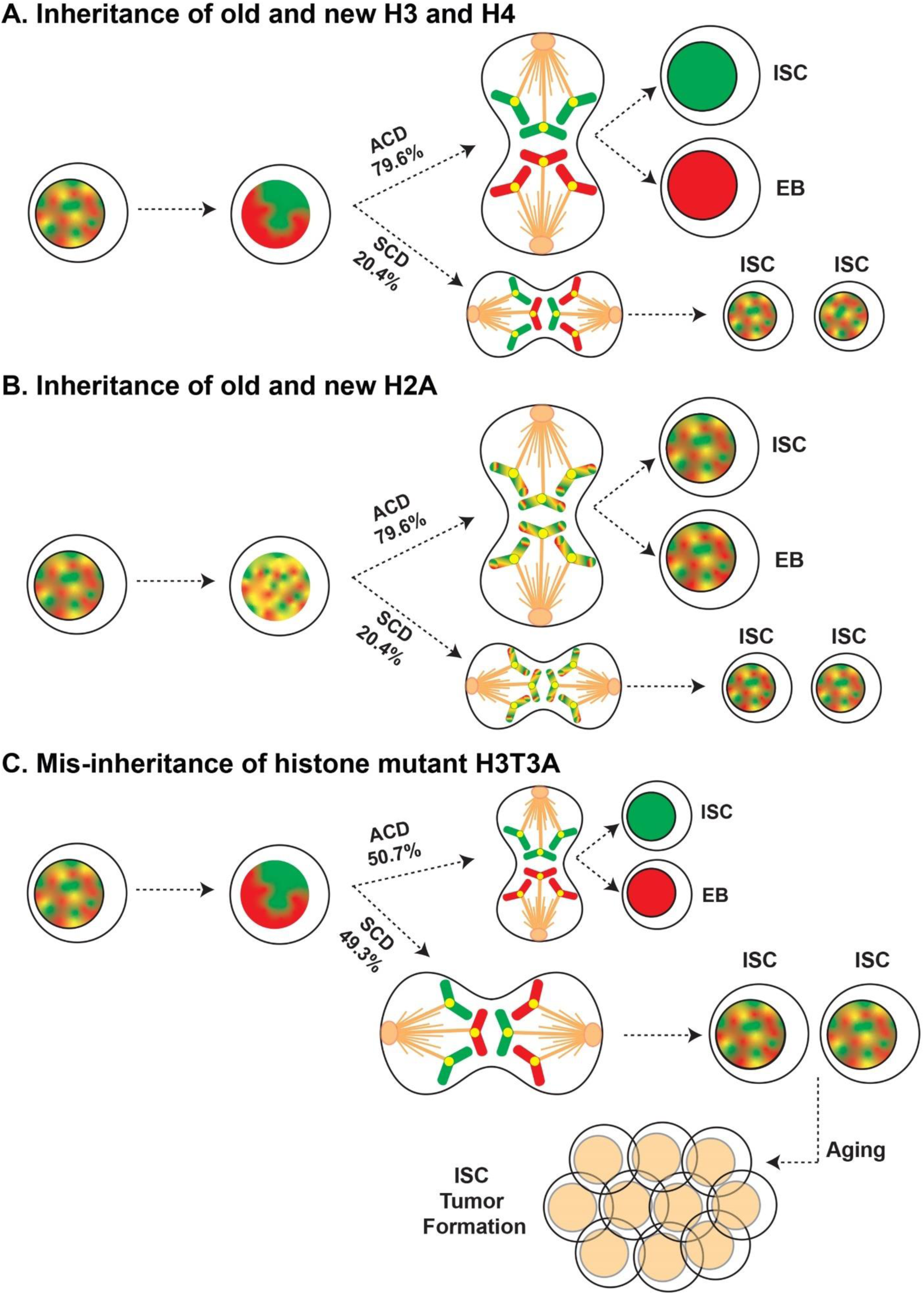
A model for different histone inheritance patterns and their roles in regulating cell identity. (**A**) Old *versus* new H3 and H4 histones are likely incorporated differentially during S phase and form separable domains visualized early in mitosis. During ACD, old *versus* new H3 and H4 enriched sister chromatids are segregated asymmetrically, giving rise to a self-renewed ISC that inherits more old H3 and H4, and an EB that inherits more new H3 and H4. During SCD, old *versus* new H3 and H4 enriched sister chromatids are segregated symmetrically, giving rise to two ISCs that inherit a comparable amount of old *versus* new H3 and H4. (**B**) Old *versus* new H2A histones are likely incorporated evenly during S phase and show an overlapping distribution visualized early in mitosis. In both ACD and SCD, old *versus* new H2A are segregated symmetrically to the resulting daughter cells. (**C**) With the mutant histone H3T3A, there is a significant increase in SCD pattern when compared to wild type H3 shown in (**A**). With aging, the mutant H3T3A also causes the formation of ISC tumors.

The biological significance of the asymmetric histone inheritance is further exemplified when this pattern is disrupted with the H3T3A histone mutant, where we detected increased SCD of ISCs as well as the formation of ISC-like tumors during aging (Figure 6C). These data connect mis-regulated histone inheritance with failures in cell fate determination and tumorigenesis. Recently, large-scale analysis identified histone mutations in 3.8% of human tumor samples, a ratio similar to the mutations of known cancer-associated genes such as *BRCA2* and *NOTCH1*. In particular, mutations at the Thr3 residue of H3 have been found in a variety of human tumor samples, including lung, breast, skin, bladder, and liver cancers (Nacev et al., 2019). However, the molecular mechanisms underlying these ‘oncohistones’ are not fully understood. Here, our findings on the oncohistone H3T3A illuminate how this mutation can lead to loss of proper epigenetic inheritance, failure in properly establishing cell identities, and the onset of tumor formation, providing a mechanistic link between mis-regulation of epigenetic inheritance and tumorigenesis.

The asymmetric inheritance mode of histones was first reported during the ACD of male *Drosophila* GSCs (Tran et al., 2012), which opened a new avenue of research; however, many essential questions remained, such as whether this asymmetric histone inheritance mode is specific to germ cells, stem cells, and/or asymmetrically dividing cells. The results reported in this paper provide a solid basis to addressing these questions. Despite significant differences in the niche structure, signaling cascades for regulating stem cell activity, and cellular differentiation pathways between the ISC and GSC lineages [(Amoyel et al., 2014; Doupe et al., 2018; Jiang et al., 2011; Jiang et al., 2009; Kiger et al., 2001; Leatherman and Dinardo, 2008; Li et al., 2014; Lin et al., 2008; Ohlstein and Spradling, 2007; Stine et al., 2014; Tulina and Matunis, 2001) and reviewed in (de Cuevas and Matunis, 2011; Kahney et al., 2019; Losick et al., 2011; Morrison and Spradling, 2008)], many features of asymmetric histone inheritance are common between these two stem cell systems. First, the cellular specificity in the GSC lineage has been shown by asymmetric histone inheritance in asymmetrically dividing GSCs but not in symmetrically dividing spermatogonial cells. In the ISC lineage, this cellular specificity is manifested by ISCs displaying an asymmetric histone inheritance mode during ACD but not during SCD. Second, this asymmetry has the molecular specificity with H3 and H4 histones in both systems, emphasizing the importance of these two canonical histones in carrying and passing on an “epigenetic memory”. Finally, expression of the mutant histone H3T3A abolishes asymmetric histone inheritance modes in both systems, resulting in stem cell or progenitor cell tumors. Therefore, these results demonstrate that the asymmetric histone inheritance mode is not specific to either germ cells or stem cells, but likely contingent on the ACD mode of mitosis with the mission to generate two distinct daughter cells, which is crucial to development, tissue homeostasis and regeneration of multicellular organisms.

Finally, it has been debated whether the two cells resulting from ISC division are intrinsically asymmetric, or only become asymmetric through the extrinsic signaling pathway after ISC division. Previous studies demonstrate that intrinsic polarity mechanisms result in the asymmetric distribution and inheritance of Par proteins to the apical daughter cell during ACD of ISCs, in order to promote differentiation (Goulas et al., 2012). Furthermore, differential Notch activities due to the Par complex activity induce distinct cellular differentiation pathways (Guo and Ohlstein, 2015). Recent work has also shown that the spindle orientation in ISCs is tightly linked with cell fate, where planar orientation gives rise to two ISCs and angular orientation generates a ISC-EB pair (Hu and Jasper, 2019). Here, through analyzing different mitotic phases of ISCs, separable old *versus* new H3 distribution are detectable in prophase and prometaphase ISCs, indicating that this asymmetry is intrinsically established prior to mitosis (Figure 6A).

Interestingly, old *versus* new H3T3A signals are also separable in prophase and prometaphase ISCs, similar to that of wild type H3. Meanwhile, increased symmetric segregation patterns in anaphase and telophase indicate that sister chromatids differentially enriched with old *versus* new H3T3A signals cannot be properly recognized and segregated. Because flies have two major autosomes (2^nd^ and 3^rd^ chromosomes) in addition to the sex chromosomes, even the randomized segregation of sister chromatids could lead to an asymmetric pattern at a low percentage, as shown previously (Xie et al., 2015). When asymmetric histone inheritance occurs with randomized segregation, the daughter cell enriched with old histone still takes the ISC cell fate (Figure 6C). Together, these findings demonstrate that ISC cell fate is likely specified by intrinsic epigenetic information and the extrinsic cues possibly ensure the differential segregation pattern. In summary, given the unique features of the ISC system, such as the ability to precisely label each derivative cell in the entire lineage, the large number of ISCs in their endogenous niche, and the sensitivity of ISC activity to environmental changes such as nutrition and aging, it will become a new *in vivo* model system to study the fundamental principles of different histone inheritance modes and relevant biological consequences under physiological and pathological conditions.

## Acknowledgements

We thank Drs. A. Spradling, W. Ludington, and the Chen laboratory members for their critical comments and suggestions on this work.

## Author Contributions

E.Z. and X.C. conceptualized the study. E.Z. performed all the experiments and data analyses. E.Z. and X.C. wrote the manuscript.

