## Supplemental Information for "Asymmetric histone inheritance regulates stem cell fate in *Drosophila* midgut"

**Materials and Methods**

**Generation of switchable dual-color transgenes.** Standard procedures were used for all molecular cloning experiments. Enzymes used for plasmid construction were obtained from New England Biolabs (Beverly, MA). The new histone sequences, such as *histone H4-mCherry* were recovered as an XbaI flanked fragment and were subsequently inserted into the XbaI site of the UASp plasmid. The old histone sequences, such as *histone H4-eGFP* were inserted to *pBluescript-FRT-NheI-SV40 PolyA-FRT* plasmid at the unique NheI site. The entire *NotI-FRT-H4-eGFP-SV40 PolyA-FRT-EcoRI* sequences were then subcloned into the *UASp-H4-mCherry* plasmid digested by *NotI* and *EcoRI*. The final *UASp-FRT-H4-eGFP-PolyA-FRT-H4-mCherry* plasmids were introduced to $w^{1118}$ flies by P-element-mediated germline transformation (Bestgene, Inc.).

**Fly strains and husbandry.** Fly stocks were raised using standard Bloomington medium at 25°C or 29°C as noted. The following fly stocks were used: *hs-flp* on the X chromosome (Bloomington Stock Center BL-26902), *esg-Gal4* on the 2$^{nd}$ chromosome (Dr. Allan Spradling, Carnegie Institute, Baltimore, MD), *Delta-nuclearLacZ* reporter on the 3$^{rd}$ chromosome (Dr. Allan Spradling, Carnegie Institute, Baltimore, MD), *UASp-FRT-H3-eGFP-PolyA-FRT-H3-mCherry* on the 2$^{nd}$ chromosome (Wooten et al., 2019), *UASp-FRT-H4-eGFP-PolyA-FRT-H4-mCherry* on the 2$^{nd}$ chromosome (this study), *UASp-FRT-H2A-eGFP-PolyA-FRT-H2A-mCherry* on the 3$^{rd}$ chromosome (Wooten et al., 2019), and *UASp-FRT-H3T3A-GFP-PolyA-FRT-H3T3A-mKO* on the 3$^{rd}$ chromosome (Xie et al., 2015).

**Heat-shock scheme.** Flies with *UASp*-dual color histone transgenes were paired with the *esg-Gal4* driver. Flies were raised at 25°C throughout development until adulthood. At eclosion, flies were transferred to new vials and aged for 10 days. On the 10$^{th}$ day, flies were transferred to new vials. Vials were then submerged underneath the water up to the plug in a circulating 37°C water bath for 90 minutes and were then recovered at 29°C for 18 hours before dissection for immunostaining experiments.

**Immunostaining experiments.** Immunofluorescence staining was performed using standard procedures (Marianes and Spradling, 2013; Tran et al., 2012). For immunofluorescence staining, midguts from 10-day old female flies were dissected in Schneider's insect media. For samples that required nucleoside analog incorporation, they were added to 10 µM EdU analog (Invitrogen Click-iT EdU Imaging Kit, catalog #C10340) and incubated for 30 minutes. All samples were then transferred to a 4% formaldehyde in phosphate-buffered saline (PBS) for 1 hour at room temperature for fixation. Samples were washed three times for five minutes in PBST (PBS with 0.1% Triton X-100), and then placed in a primary antibody solution containing antibodies at desired concentrations in 5% normal goat serum (NGS) in PBST. Samples were incubated for a minimum of 24 hours at 4°C in primary antibodies. Antibodies used in these experiments are mouse anti-H3S10ph (1:1000, Abcam AB-14995) and chicken anti-ß-galactosidase (1:1000, Abcam AB-9361). All GFP, mCherry, and mKO signal was visualized using endogenous fluorescence. Samples were then wash three times, five minutes each time in PBST, and then incubated in a 1:1000 dilution of Alexa-Flour-conjugated secondary antibody in 5% NGS in PBST for 2 hours at room temperature. For EdU visualization, EdU analog was conjugated to Alexa-647 dye using CLICK chemistry [reviewed in (Kolb et al., 2001; Nwe and Brechbiel,

2009)]. Samples were washed three times, five minutes each time in PBST, and then mounted for microscopy in Vectashield antifade mounting medium (Vector Laboratories, Cat#H-1400) with or without DAPI. Samples were imaged using a Leica SP8 confocal microscope with a 63x oil immersion objective. Images were analyzed using ImageJ software.

**Quantification of fluorescent signals.** For all quantification, GFP, mCherry, and mKO signals were not enhanced with antibody, and the endogenous fluorescence was used. Values of GFP, mCherry, mKO, and Dl-nLacZ were calculated using ImageJ software. The cell or region of interest was outlined, and the raw integrated density was recorded for each channel of interest. A background measurement was then taken using the same outlined region that was measured in an area with no fluorescence signal. The background measurement was subtracted from the initial measurement. For every cell, each z slice containing that cell was included for quantification, and the values were added together to get the total value. These values were then used to compute the ratios of interest.

**Quantification of histone asymmetry in post-mitotic pairs.** To quantify the ratios of old and new histones in post-mitotic pairs, the pairs were first selected using the following criteria. The two cells must be within 5 μm of each other, one cell must be no more than 1.5X larger than the other cell based on cell diameter to avoid using the polyploid enterocytes (EC), both cells must be EdU negative, and both cells must contain quantifiable levels of old and new histone. Quantification of old histone, new histone, and Dl-nLacZ was conducted using the method described above for each cell. The cell with a higher Dl-nLacZ value was denoted as cell 1, and the other as cell 2. A ratio was computed by dividing the values of old histone, new histone, and

Dl-nLacZ for cell 1 by cell 2. Any pair with a Dl-nLacZ ratio greater than 2 was considered an asymmetric pair, while any pair with a ratio less than 2 was considered a symmetric pair. The ratios for old and new histone were then recorded as either a ratio of ISC/EB or ISC1/ISC2, where ISC1 had the relatively higher Dl-nLacZ value compared to ISC2.

**Quantification of histone asymmetry in anaphase and telophase mitotic cells.** Anaphase and telophase cells were identified based on the mitotic marker H3S10ph and chromosomal morphology. The sister chromatids were required to be segregated far enough to show separation, distinguishing the two individual sets. Each set of sister chromatids was outlined, and old histone was measured using the method described above. A ratio was computed by dividing the sister chromatids with the higher value of old histone by sister chromatids with the lower value of old histone.

**Correlation analysis between old and new histones in prophase and prometaphase cells.** Prophase and prometaphase cells were identified based on the mitotic marker H3S10ph and morphology. The center of this cell was identified and outlined as a region of interest, and the image was split into two channels, old histone (eGFP at 488nm) and new histone (mCherry at 568nm), for image analysis. The Coloc2 analysis tool in ImageJ was used to collect the Pearson and Spearman correlation coefficients. The old and new histone channels were compared, and the Pearson and Spearman correlation coefficients were recorded.

**Statistics and reproducibility.** Data was subjected to the Shapiro-Wilk test to determine whether the data was normally distributed or skewed. For normally distributed data, a single sample t test was used for comparing one dataset to a hypothesized mean of 0 or 1. For skewed data, the Wilcoxon signed rank test was used to compare one dataset to a hypothesized median of 0 or 1. An unpaired two sample t test was used to compare two individual datasets to each other. Data are presented with error bars representing the mean $\pm$ SE (standard error). Significant differences based on these statistical analyses were noted by asterisks (* $p < 0.05$, ** $p < 0.01$, *** $p < 0.001$. **** $p < 0.0001$).

**Supplemental Figures and Figure Legends:**

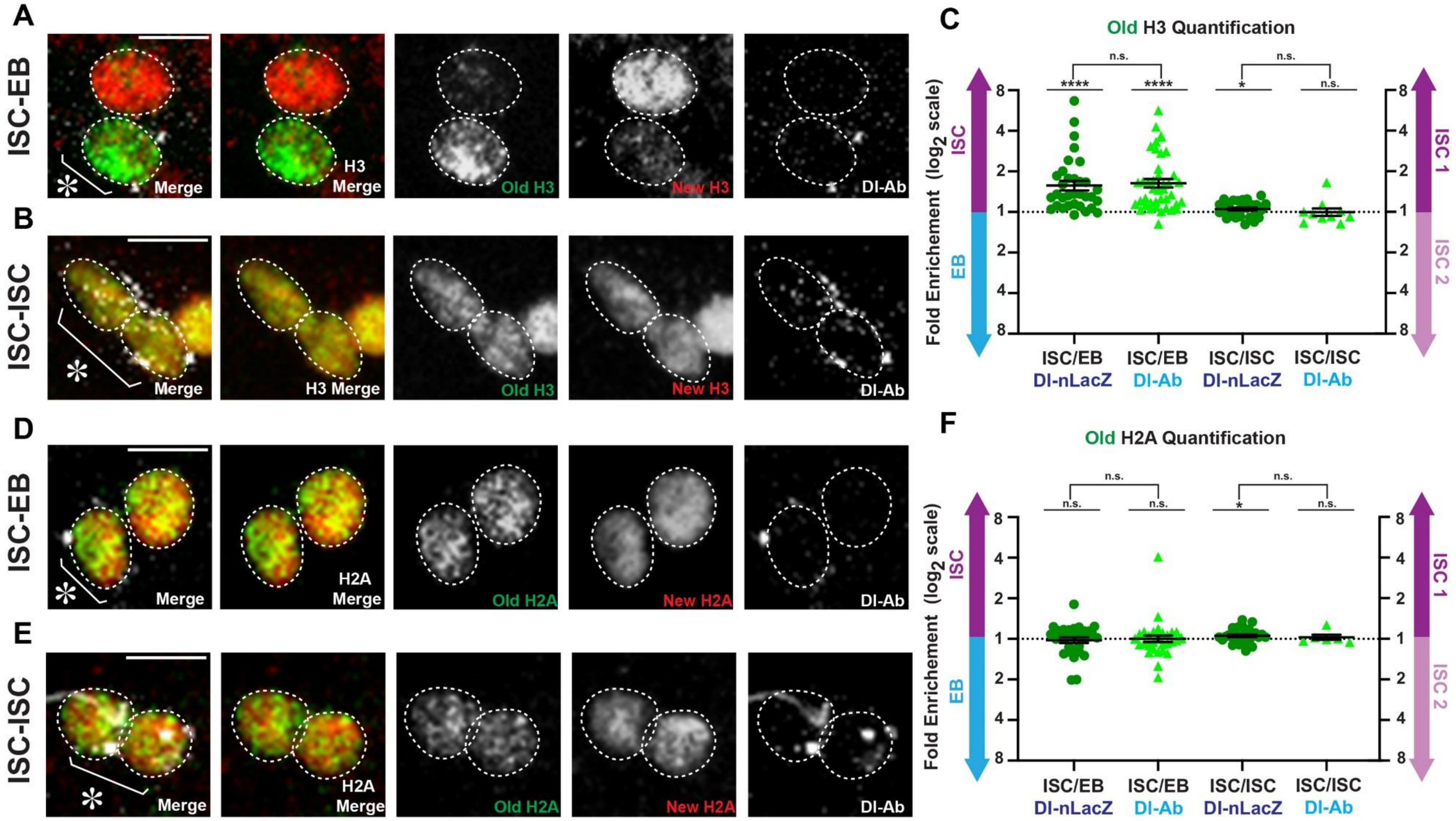

**Figure S1: Old and new histone H3 display asymmetric inheritance mode specific for asymmetric division of intestinal stem cells using Delta antibody.** (**A**) Old and new H3 distribution in a post-mitotic ISC-EB pair with punctate Delta antibody staining to identify the ISC, showing that H3-eGFP (old) is asymmetrically inherited by the ISC, while H3-mCherry (new) is enriched in the EB. (**B**) Old and new H3 distribution in a post-mitotic ISC-ISC pair with punctate Delta antibody staining to identify ISCs, showing that both H3-eGFP (old) and H3-mCherry (new) are symmetrically distributed between the two ISC nuclei. (**C**) Comparisons of the quantification of old H3 in ISC-EB and ISC-ISC pairs using the Delta-nLacZ reporter line (data from Figure 1F), and the Delta antibody to identify ISCs. Quantification of H3-eGFP (old) in ISC-EB pairs identified by the Delta antibody (avg. $\log_2$ ratio for old H3 = 0.71 $\pm$ 0.10, n = 41) is similar to that of ISC-EB pairs identified by the Delta-nLacZ reporter (avg. $\log_2$ ratio for old H3 = 0.65 $\pm$ 0.12, n = 33, Figure 1F). Similarly, quantification of H3-eGFP (old) in ISC-ISC pairs was similar between pairs identified by the Delta antibody (avg. $\log_2$ ratio for old H3 = -0.003 $\pm$ 0.09, n = 10) and pairs identified by the Delta-nLacZ reporter (avg. $\log_2$ ratio for old H3 = 0.07 $\pm$ 0.03, n = 30, Figure 1F). (**D**) Old and new H2A distribution in a post-mitotic ISC-EB pair with punctate Delta antibody staining to identify the ISC, showing that H2A-eGFP (old) and H2A-mCherry (new) are symmetrically inherited in the ISC and EB. (**E**) Old and new H2A distribution in a post-mitotic ISC-ISC pair with punctate Delta antibody staining to identify ISCs, showing that both H2A-eGFP (old) and H2A-mCherry (new) are symmetrically distributed between the two nuclei. (**F**) Comparisons of the quantification of old H2A in ISC-EB and ISC-ISC pairs using the Delta-nLacZ reporter line (data from Figure 2F), or the Delta antibody to identify ISCs. Quantification of H2A-eGFP (old) in ISC-EB pairs identified by the Delta antibody (avg. $\log_2$ ratio for old H2A = 0.002 $\pm$ 0.08, n = 32) is similar to that of ISC-EB pairs identified by the Delta-nLacZ reporter (avg. $\log_2$ ratio for old H2A = -0.03 $\pm$ 0.07, n = 30, Figure 2F). Similarly, quantification of H2A-eGFP (old) in ISC-ISC pairs was similar between pairs identified by the Delta antibody (avg. $\log_2$ ratio for old H2A = 0.03 $\pm$ 0.06, n = 6) and pairs identified by the Delta-nLacZ reporter (avg. $\log_2$ ratio for old H2A = 0.08 $\pm$ 0.03, n = 30, Figure 2F). For (**C**) and (**F**), individual data points and mean values are shown. Error bars represent SEM. **** $p < 0.0001$, * $p < 0.05$; single sample t test (for normally distributed data) for comparing one dataset to a hypothesized mean of 0 ($\log_2$ value representing a 1:1 ratio), or Wilcoxon signed rank test (for skewed data) for comparing one dataset to a hypothesized median of 0. Unpaired t test to compare two individual datasets to each other. NS, not significant. Individual data values are shown in Supplemental Table 1. Scale bar in (**A**), (**B**), (**D**) and (**E**): 5 µm; asterisk, ISC side.

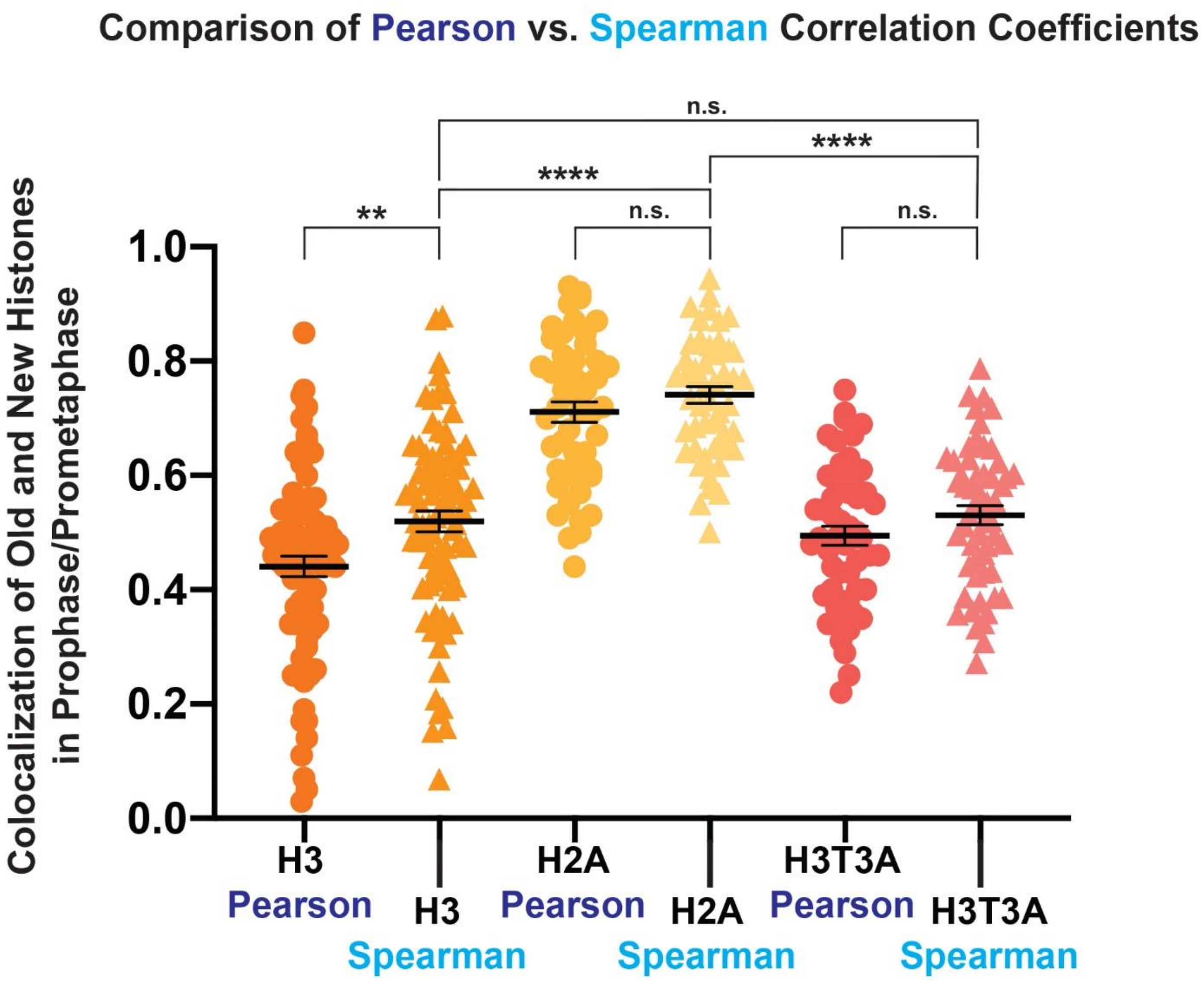

**Figure S2: Comparison of Pearson *versus* Spearman correlation coefficients in determining the colocalization of old *versus* new histones for H3, H2A, and H3T3A in prophase and prometaphase ISCs.** Comparison of Pearson and Spearman correlation coefficients to determine the colocalization of old *versus* new histones in prophase and prometaphase ISCs. Spearman correlation coefficients are slightly higher than Pearson correlation coefficients. There are no significant differences between Pearson and Spearman correlation coefficients for H2A [avg. Pearson correlation coefficient for H2A = 0.71 $\pm$ 0.02 (Figure 3G), avg. Spearman correlation coefficient for H2A = 0.74 $\pm$ 0.01, n = 50] and H3T3A [avg. Pearson correlation coefficient for H3T3A = 0.49 $\pm$ 0.02 (Figure 4F), avg. Spearman correlation coefficient for H3T3A = 0.53 $\pm$ 0.02, n = 55] analyses. There is a significant difference between the Pearson and Spearman correlation coefficients for H3 [avg. Pearson correlation coefficient for H3 = 0.44 $\pm$ 0.02 (Figure 3G), avg. Spearman correlation coefficient for H3 = 0.51 $\pm$ 0.02, n = 50, ** $p < 0.01$); however, the general trend of correlation coefficients is maintained between the two methods of analysis. Spearman's analysis demonstrates that H2A correlation coefficients are significantly higher than H3 and H3T3A (**** $p < 0.0001$), while there is no significant difference between H3 and H3T3A correlation coefficients (n.s.). This is consistent with the results of Pearson's analysis (shown in Figures 3G and 4F). Individual data points and mean values are shown. Error bars represent SEM. **** $p < 0.0001$, ** $p < 0.01$; unpaired t test to compare two individual datasets to each other. NS, not significant. Individual data values are shown in Supplemental Table 3.

**Supplementary Tables:**

**Supplemental Table 1: Quantification of old and new histone H3, H4, and H2A in post-mitotic pairs identified by Delta-nLacZ or Delta antibody with imaging on fixed samples.**

*H3 Data with Delta-nLacZ:*

| Pair # | Old ISC/EB | Log2 (old) | New ISC/EB | Log2 (new) | Pair # | Old ISC1/ISC2 | Log2 (old) | New ISC1/ISC2 | Log2 (new) |
|---|---|---|---|---|---|---|---|---|---|
| 1 | 3.662 | 1.873 | 0.623 | -0.684 | 1 | 0.986 | -0.020 | 1.015 | 0.022 |
| 2 | 1.281 | 0.357 | 0.386 | -1.373 | 2 | 0.996 | -0.006 | 1.153 | 0.205 |
| 3 | 1.049 | 0.069 | 0.848 | -0.238 | 3 | 1.086 | 0.118 | 1.051 | 0.071 |
| 4 | 1.680 | 0.748 | 0.263 | -1.927 | 4 | 0.906 | -0.143 | 1.110 | 0.150 |
| 5 | 4.645 | 2.216 | 1.910 | 0.934 | 5 | 1.221 | 0.288 | 1.538 | 0.621 |
| 6 | 1.765 | 0.820 | 0.492 | -1.022 | 6 | 1.022 | 0.031 | 0.868 | -0.205 |
| 7 | 1.209 | 0.274 | 0.755 | -0.406 | 7 | 0.893 | -0.163 | 0.792 | -0.336 |
| 8 | 2.014 | 1.010 | 1.344 | 0.427 | 8 | 0.907 | -0.140 | 1.029 | 0.041 |
| 9 | 1.668 | 0.738 | 0.264 | -1.922 | 9 | 0.911 | -0.135 | 0.921 | -0.118 |
| 10 | 2.998 | 1.584 | 0.691 | -0.534 | 10 | 1.063 | 0.088 | 1.055 | 0.078 |
| 11 | 1.597 | 0.676 | 0.906 | -0.142 | 11 | 1.136 | 0.184 | 1.026 | 0.037 |
| 12 | 6.667 | 2.737 | 0.380 | -1.395 | 12 | 1.131 | 0.178 | 1.029 | 0.041 |
| 13 | 1.111 | 0.152 | 1.079 | 0.110 | 13 | 1.136 | 0.183 | 1.243 | 0.314 |
| 14 | 2.370 | 1.245 | 1.163 | 0.218 | 14 | 1.042 | 0.059 | 1.024 | 0.034 |
| 15 | 1.145 | 0.196 | 0.991 | -0.013 | 15 | 0.934 | -0.099 | 0.964 | -0.052 |
| 16 | 1.278 | 0.354 | 0.683 | -0.549 | 16 | 0.808 | -0.308 | 1.042 | 0.059 |
| 17 | 1.103 | 0.142 | 0.731 | -0.453 | 17 | 1.074 | 0.102 | 1.032 | 0.045 |
| 18 | 0.989 | -0.017 | 0.648 | -0.627 | 18 | 1.247 | 0.318 | 1.319 | 0.399 |
| 19 | 1.294 | 0.372 | 1.006 | 0.009 | 19 | 1.131 | 0.178 | 1.080 | 0.111 |
| 20 | 1.358 | 0.441 | 0.383 | -1.384 | 20 | 1.232 | 0.301 | 1.040 | 0.056 |
| 21 | 1.066 | 0.093 | 0.421 | -1.247 | 21 | 0.843 | -0.246 | 0.901 | -0.150 |
| 22 | 1.357 | 0.440 | 0.655 | -0.611 | 22 | 0.924 | -0.114 | 0.819 | -0.288 |
| 23 | 2.414 | 1.271 | 1.446 | 0.532 | 23 | 0.993 | -0.011 | 1.011 | 0.016 |
| 24 | 1.777 | 0.830 | 0.896 | -0.158 | 24 | 1.118 | 0.160 | 0.997 | -0.005 |
| 25 | 1.111 | 0.152 | 0.560 | -0.837 | 25 | 1.244 | 0.316 | 1.041 | 0.058 |
| 26 | 1.353 | 0.436 | 0.333 | -1.585 | 26 | 1.116 | 0.158 | 1.097 | 0.134 |
| 27 | 1.459 | 0.545 | 1.094 | 0.130 | 27 | 1.325 | 0.406 | 0.905 | -0.144 |
| 28 | 1.005 | 0.008 | 0.520 | -0.944 | 28 | 1.213 | 0.278 | 0.964 | -0.053 |
| 29 | 0.951 | -0.073 | 0.699 | -0.516 | 29 | 1.147 | 0.198 | 0.973 | -0.039 |
| 30 | 1.854 | 0.890 | 0.891 | -0.167 | 30 | 1.032 | 0.045 | 0.929 | -0.106 |
| 31 | 1.328 | 0.410 | 0.800 | -0.322 | | | | | |
| 32 | 1.066 | 0.093 | 0.461 | -1.117 | | | | | |
| 33 | 1.299 | 0.377 | 0.578 | -0.792 | | | | | |

*H4 Data with Delta-nLacZ:*

| Pair # | Old ISC/EB | Log2 (old) | New ISC/EB | Log2 (new) | Pair # | Old ISC1/ISC2 | Log2 (old) | New ISC1/ISC2 | Log2 (new) |
|---|---|---|---|---|---|---|---|---|---|
| 1 | 0.911 | -0.135 | 0.693 | -0.529 | 1 | 0.965 | -0.052 | 1.026 | 0.038 |
| 2 | 1.988 | 0.991 | 0.751 | -0.413 | 2 | 0.897 | -0.157 | 1.211 | 0.277 |
| 3 | 1.090 | 0.124 | 0.244 | -2.038 | 3 | 0.981 | -0.028 | 0.975 | -0.036 |
| 4 | 1.178 | 0.236 | 0.465 | -1.105 | 4 | 0.990 | -0.014 | 0.822 | -0.282 |
| 5 | 1.093 | 0.128 | 0.790 | -0.340 | 5 | 1.063 | 0.088 | 1.146 | 0.197 |
| 6 | 2.293 | 1.197 | 0.573 | -0.804 | 6 | 1.095 | 0.131 | 1.282 | 0.359 |
| 7 | 1.140 | 0.189 | 0.909 | -0.138 | 7 | 0.837 | -0.257 | 0.664 | -0.591 |
| 8 | 1.087 | 0.120 | 0.693 | -0.530 | 8 | 1.131 | 0.178 | 0.978 | -0.031 |
| 9 | 1.124 | 0.169 | 0.898 | -0.154 | 9 | 1.100 | 0.137 | 1.129 | 0.175 |
| 10 | 1.750 | 0.807 | 1.086 | 0.119 | 10 | 1.045 | 0.064 | 0.931 | -0.104 |
| 11 | 1.116 | 0.158 | 1.366 | 0.450 | 11 | 1.085 | 0.118 | 0.973 | -0.039 |
| 12 | 1.130 | 0.177 | 0.639 | -0.645 | 12 | 0.898 | -0.155 | 0.990 | -0.014 |
| 13 | 1.154 | 0.207 | 0.627 | -0.673 | 13 | 1.001 | 0.001 | 0.658 | -0.604 |
| 14 | 1.597 | 0.675 | 0.788 | -0.344 | 14 | 0.933 | -0.100 | 0.874 | -0.194 |
| 15 | 1.691 | 0.758 | 0.692 | -0.531 | 15 | 1.035 | 0.050 | 0.840 | -0.252 |
| 16 | 1.628 | 0.703 | 0.782 | -0.355 | 16 | 0.955 | -0.066 | 1.036 | 0.051 |
| 17 | 2.114 | 1.080 | 0.809 | -0.305 | 17 | 0.833 | -0.263 | 0.831 | -0.267 |
| 18 | 1.237 | 0.306 | 0.479 | -1.063 | 18 | 1.054 | 0.075 | 0.943 | -0.084 |
| 19 | 1.561 | 0.643 | 0.992 | -0.011 | 19 | 1.193 | 0.255 | 0.847 | -0.240 |

*H2A Data with Delta-nLacZ:*

| Pair # | Old ISC/EB | Log2 (old) | New ISC/EB | Log2 (new) | Pair # | Old ISC1/ISC2 | Log2 (old) | New ISC1/ISC2 | Log2 (new) |
|---|---|---|---|---|---|---|---|---|---|
| 1 | 1.080 | 0.111 | 0.727 | -0.461 | 1 | 1.232 | 0.301 | 1.020 | 0.029 |
| 2 | 0.881 | -0.183 | 1.101 | 0.139 | 2 | 0.895 | -0.160 | 0.654 | -0.614 |
| 3 | 1.808 | 0.854 | 1.617 | 0.694 | 3 | 1.147 | 0.198 | 0.995 | -0.008 |
| 4 | 1.222 | 0.289 | 1.577 | 0.657 | 4 | 0.986 | -0.021 | 0.867 | -0.205 |
| 5 | 0.867 | -0.206 | 0.787 | -0.345 | 5 | 0.872 | -0.197 | 0.835 | -0.259 |
| 6 | 1.040 | 0.056 | 1.115 | 0.158 | 6 | 1.029 | 0.041 | 0.841 | -0.250 |
| 7 | 1.022 | 0.032 | 0.916 | -0.126 | 7 | 1.203 | 0.267 | 1.216 | 0.283 |
| 8 | 0.774 | -0.370 | 0.565 | -0.823 | 8 | 0.943 | -0.084 | 0.676 | -0.564 |
| 9 | 0.801 | -0.320 | 0.607 | -0.721 | 9 | 1.041 | 0.058 | 1.055 | 0.078 |
| 10 | 1.164 | 0.219 | 0.962 | -0.056 | 10 | 1.248 | 0.320 | 0.904 | -0.146 |
| 11 | 0.751 | -0.413 | 0.988 | -0.018 | 11 | 1.093 | 0.128 | 0.653 | -0.614 |
| 12 | 0.833 | -0.263 | 1.777 | 0.829 | 12 | 1.022 | 0.031 | 0.684 | -0.548 |
| 13 | 0.496 | -1.011 | 0.536 | -0.901 | 13 | 1.071 | 0.099 | 0.931 | -0.103 |
| 14 | 0.990 | -0.015 | 0.480 | -1.060 | 14 | 1.031 | 0.044 | 1.013 | 0.019 |
| 15 | 1.151 | 0.203 | 1.337 | 0.419 | 15 | 1.175 | 0.232 | 1.000 | 0.000 |
| 16 | 1.169 | 0.225 | 0.773 | -0.372 | 16 | 1.039 | 0.055 | 0.754 | -0.408 |
| 17 | 1.025 | 0.035 | 0.767 | -0.383 | 17 | 1.384 | 0.468 | 0.918 | -0.124 |

| | | | | | | | | | |
|---|---|---|---|---|---|---|---|---|---|
| 18 | 0.499 | -1.004 | 1.198 | 0.261 | 18 | 1.029 | 0.042 | 1.012 | 0.017 |
| 19 | 1.101 | 0.139 | 0.770 | -0.376 | 19 | 1.037 | 0.053 | 0.952 | -0.071 |
| 20 | 1.111 | 0.152 | 0.775 | -0.367 | 20 | 0.813 | -0.298 | 0.926 | -0.111 |
| 21 | 1.142 | 0.191 | 0.639 | -0.647 | 21 | 1.338 | 0.420 | 1.369 | 0.453 |
| 22 | 1.192 | 0.254 | 1.256 | 0.329 | 22 | 0.955 | -0.066 | 0.978 | -0.032 |
| 23 | 1.227 | 0.295 | 0.945 | -0.082 | 23 | 0.967 | -0.048 | 0.899 | -0.153 |
| 24 | 1.018 | 0.026 | 0.972 | -0.041 | 24 | 1.104 | 0.142 | 0.977 | -0.034 |
| 25 | 1.236 | 0.306 | 0.959 | -0.061 | 25 | 0.905 | -0.144 | 1.005 | 0.008 |
| 26 | 0.798 | -0.325 | 0.461 | -1.119 | 26 | 1.077 | 0.107 | 1.260 | 0.333 |
| 27 | 0.998 | -0.002 | 1.191 | 0.252 | 27 | 0.944 | -0.083 | 1.043 | 0.060 |
| 28 | 1.144 | 0.194 | 0.910 | -0.136 | 28 | 1.050 | 0.071 | 1.194 | 0.255 |
| 29 | 1.008 | 0.011 | 0.916 | -0.127 | 29 | 1.029 | 0.041 | 0.839 | -0.254 |
| 30 | 0.728 | -0.459 | 0.928 | -0.107 | 30 | 1.209 | 0.274 | 1.103 | 0.141 |

*H3 Data with Delta Antibody:*

| Pair # | Old ISC/EB | Log2 (old) | Pair # | Old ISC1/ISC2 | Log2 (old) |
|---|---|---|---|---|---|
| 1 | 4.281 | 2.098 | 1 | 0.921 | -0.118 |
| 2 | 1.060 | 0.084 | 2 | 0.904 | -0.145 |
| 3 | 0.816 | -0.293 | 3 | 1.659 | 0.730 |
| 4 | 1.248 | 0.320 | 4 | 0.819 | -0.288 |
| 5 | 5.667 | 2.503 | 5 | 0.831 | -0.267 |
| 6 | 1.010 | 0.014 | 6 | 1.005 | 0.007 |
| 7 | 2.737 | 1.452 | 7 | 1.130 | 0.176 |
| 8 | 1.070 | 0.097 | 8 | 1.018 | 0.025 |
| 9 | 3.073 | 1.620 | 9 | 0.972 | -0.042 |
| 10 | 1.192 | 0.254 | 10 | 0.927 | -0.110 |
| 11 | 1.872 | 0.905 | | | |
| 12 | 2.092 | 1.065 | | | |
| 13 | 1.034 | 0.048 | | | |
| 14 | 1.209 | 0.274 | | | |
| 15 | 1.657 | 0.728 | | | |
| 16 | 3.599 | 1.847 | | | |
| 17 | 1.494 | 0.580 | | | |
| 18 | 2.812 | 1.491 | | | |
| 19 | 3.324 | 1.733 | | | |
| 20 | 1.177 | 0.235 | | | |
| 21 | 1.190 | 0.251 | | | |
| 22 | 1.036 | 0.051 | | | |
| 23 | 1.489 | 0.574 | | | |
| 24 | 3.678 | 1.879 | | | |
| 25 | 1.714 | 0.777 | | | |
| 26 | 1.860 | 0.896 | | | |
| 27 | 1.144 | 0.194 | | | |

| | | |
|---|---|---|
| 28 | 1.660 | 0.731 |
| 29 | 1.122 | 0.166 |
| 30 | 1.137 | 0.185 |
| 31 | 1.340 | 0.422 |
| 32 | 1.056 | 0.079 |
| 33 | 1.607 | 0.684 |
| 34 | 1.671 | 0.740 |
| 35 | 1.684 | 0.752 |
| 36 | 2.945 | 1.558 |
| 37 | 1.254 | 0.326 |
| 38 | 1.094 | 0.129 |
| 39 | 1.627 | 0.702 |
| 40 | 1.459 | 0.545 |
| 41 | 1.265 | 0.339 |

*H2A Data with Delta Antibody:*

| Pair # | Old ISC/EB | Log2 (old) | Pair # | Old ISC1/ISC2 | Log2 (old) |
|---|---|---|---|---|---|
| 1 | 0.821 | -0.285 | 1 | 1.003 | 0.005 |
| 2 | 1.069 | 0.097 | 2 | 1.270 | 0.344 |
| 3 | 0.999 | -0.002 | 3 | 1.021 | 0.031 |
| 4 | 1.130 | 0.177 | 4 | 0.944 | -0.082 |
| 5 | 0.518 | -0.948 | 5 | 0.967 | -0.048 |
| 6 | 4.067 | 2.024 | 6 | 0.991 | -0.014 |
| 7 | 1.060 | 0.085 | | | |
| 8 | 1.131 | 0.178 | | | |
| 9 | 1.110 | 0.150 | | | |
| 10 | 1.007 | 0.011 | | | |
| 11 | 1.006 | 0.008 | | | |
| 12 | 1.028 | 0.040 | | | |
| 13 | 1.046 | 0.065 | | | |
| 14 | 0.923 | -0.115 | | | |
| 15 | 0.790 | -0.340 | | | |
| 16 | 1.088 | 0.121 | | | |
| 17 | 1.459 | 0.545 | | | |
| 18 | 0.779 | -0.359 | | | |
| 19 | 0.799 | -0.323 | | | |
| 20 | 0.931 | -0.103 | | | |
| 21 | 0.945 | -0.082 | | | |
| 22 | 0.629 | -0.669 | | | |
| 23 | 0.952 | -0.071 | | | |
| 24 | 0.907 | -0.142 | | | |
| 25 | 0.971 | -0.042 | | | |
| 26 | 1.123 | 0.167 | | | |

| | | |
|---|---|---|
| 27 | 1.190 | 0.251 |
| 28 | 0.842 | -0.247 |
| 29 | 0.911 | -0.135 |
| 30 | 0.895 | -0.160 |
| 31 | 1.126 | 0.171 |
| 32 | 0.999 | -0.002 |

**Supplemental Table 2: Quantification of old histone asymmetry in anaphase and telophase mitotic cells for H3, H2A, and H3T3A**

| Pair # | Old Histone H3 Ratio | Old Histone H2A Ratio | Old Histone H3T3A Ratio |
|---|---|---|---|
| 1 | 1.20 | 1.16 | 1.15 |
| 2 | 2.20 | 1.04 | 1.08 |
| 3 | 1.67 | 1.12 | 1.08 |
| 4 | 1.48 | 1.15 | 1.02 |
| 5 | 2.63 | 1.04 | 1.04 |
| 6 | 1.73 | 1.28 | 1.06 |
| 7 | 1.49 | 1.03 | 1.16 |
| 8 | 1.11 | 1.05 | 1.15 |
| 9 | 1.20 | 1.04 | 1.32 |
| 10 | 1.52 | 1.04 | 1.07 |
| 11 | 1.23 | 1.10 | 1.06 |
| 12 | 1.33 | 1.15 | |
| 13 | 1.02 | 1.03 | |
| 14 | 1.07 | 1.02 | |
| 15 | 1.27 | 1.09 | |
| 16 | 1.38 | 1.12 | |
| 17 | 1.26 | 1.09 | |
| 18 | | 1.12 | |
| 19 | | 1.00 | |
| 20 | | 1.01 | |

**Supplemental Table 3: Quantification of colocalization between GFP and mCherry or mKO tagged histones H3, H2A, and H3T3A.**

*H3 Data:*

| Cell # | Pearson Correlation Coefficient | Spearman Correlation Coefficient |
|---|---|---|
| 1 | 0.31 | 0.469 |
| 2 | 0.72 | 0.874 |
| 3 | 0.14 | 0.185 |
| 4 | 0.51 | 0.655 |
| 5 | 0.38 | 0.578 |
| 6 | 0.34 | 0.416 |
| 7 | 0.42 | 0.632 |
| 8 | 0.46 | 0.499 |
| 9 | 0.49 | 0.584 |
| 10 | 0.64 | 0.693 |
| 11 | 0.45 | 0.547 |
| 12 | 0.34 | 0.487 |
| 13 | 0.47 | 0.593 |
| 14 | 0.19 | 0.4 |
| 15 | 0.57 | 0.539 |
| 16 | 0.05 | 0.159 |
| 17 | 0.52 | 0.534 |
| 18 | 0.42 | 0.515 |
| 19 | 0.26 | 0.403 |
| 20 | 0.34 | 0.434 |
| 21 | 0.25 | 0.428 |
| 22 | 0.55 | 0.652 |
| 23 | 0.45 | 0.567 |
| 24 | 0.47 | 0.74 |
| 25 | 0.7 | 0.746 |
| 26 | 0.66 | 0.711 |
| 27 | 0.75 | 0.798 |
| 28 | 0.51 | 0.618 |
| 29 | 0.85 | 0.879 |
| 30 | 0.47 | 0.601 |
| 31 | 0.11 | 0.194 |
| 32 | 0.25 | 0.342 |
| 33 | 0.56 | 0.629 |
| 34 | 0.48 | 0.568 |
| 35 | 0.5 | 0.637 |
| 36 | 0.74 | 0.776 |
| 37 | 0.03 | 0.069 |

| | | |
|---|---|---|
| 38 | 0.17 | 0.358 |
| 39 | 0.07 | 0.152 |
| 40 | 0.5 | 0.527 |
| 41 | 0.49 | 0.585 |
| 42 | 0.54 | 0.612 |
| 43 | 0.53 | 0.654 |
| 44 | 0.62 | 0.676 |
| 45 | 0.64 | 0.607 |
| 46 | 0.5 | 0.554 |
| 47 | 0.55 | 0.604 |
| 48 | 0.46 | 0.519 |
| 49 | 0.54 | 0.653 |
| 50 | 0.44 | 0.604 |
| 51 | 0.43 | 0.456 |
| 52 | 0.44 | 0.515 |
| 53 | 0.56 | 0.637 |
| 54 | 0.46 | 0.484 |
| 55 | 0.54 | 0.556 |
| 56 | 0.54 | 0.633 |
| 57 | 0.49 | 0.571 |
| 58 | 0.44 | 0.567 |
| 59 | 0.51 | 0.577 |
| 60 | 0.46 | 0.486 |
| 61 | 0.67 | 0.745 |
| 62 | 0.64 | 0.679 |
| 63 | 0.4 | 0.406 |
| 64 | 0.6 | 0.735 |
| 65 | 0.42 | 0.44 |
| 66 | 0.48 | 0.514 |
| 67 | 0.36 | 0.3 |
| 68 | 0.3 | 0.331 |
| 69 | 0.44 | 0.474 |
| 70 | 0.43 | 0.476 |
| 71 | 0.28 | 0.353 |
| 72 | 0.48 | 0.561 |
| 73 | 0.24 | 0.257 |
| 74 | 0.25 | 0.324 |
| 75 | 0.39 | 0.445 |
| 76 | 0.47 | 0.505 |
| 77 | 0.37 | 0.485 |
| 78 | 0.33 | 0.329 |
| 79 | 0.37 | 0.409 |
| 80 | 0.34 | 0.345 |
| 81 | 0.17 | 0.209 |

*H2A Data:*

| Cell # | Pearson Correlation Coefficient | Spearman Correlation Coefficient |
|---|---|---|
| 1 | 0.85 | 0.871 |
| 2 | 0.93 | 0.945 |
| 3 | 0.8 | 0.828 |
| 4 | 0.79 | 0.778 |
| 5 | 0.66 | 0.678 |
| 6 | 0.53 | 0.63 |
| 7 | 0.64 | 0.692 |
| 8 | 0.71 | 0.735 |
| 9 | 0.85 | 0.873 |
| 10 | 0.76 | 0.803 |
| 11 | 0.57 | 0.618 |
| 12 | 0.64 | 0.685 |
| 13 | 0.83 | 0.796 |
| 14 | 0.67 | 0.732 |
| 15 | 0.75 | 0.779 |
| 16 | 0.84 | 0.84 |
| 17 | 0.87 | 0.787 |
| 18 | 0.77 | 0.676 |
| 19 | 0.81 | 0.831 |
| 20 | 0.74 | 0.771 |
| 21 | 0.72 | 0.774 |
| 22 | 0.77 | 0.821 |
| 23 | 0.86 | 0.89 |
| 24 | 0.9 | 0.836 |
| 25 | 0.87 | 0.879 |
| 26 | 0.52 | 0.578 |
| 27 | 0.61 | 0.654 |
| 28 | 0.44 | 0.569 |
| 29 | 0.61 | 0.678 |
| 30 | 0.92 | 0.914 |
| 31 | 0.78 | 0.764 |
| 32 | 0.58 | 0.648 |
| 33 | 0.7 | 0.695 |
| 34 | 0.91 | 0.895 |
| 35 | 0.75 | 0.817 |
| 36 | 0.6 | 0.641 |
| 37 | 0.49 | 0.501 |
| 38 | 0.65 | 0.711 |
| 39 | 0.55 | 0.648 |
| 40 | 0.61 | 0.646 |

| | | |
|---|---|---|
| 41 | 0.79 | 0.819 |
| 42 | 0.77 | 0.757 |
| 43 | 0.61 | 0.724 |
| 44 | 0.6 | 0.617 |
| 45 | 0.68 | 0.769 |
| 46 | 0.5 | 0.55 |
| 47 | 0.71 | 0.775 |
| 48 | 0.53 | 0.596 |
| 49 | 0.8 | 0.813 |
| 50 | 0.72 | 0.723 |

H3T3A Data:

| **Cell #** | **Pearson Correlation Coefficient** | **Spearman Correlation Coefficient** |
|---|---|---|
| 1 | 0.59 | 0.631 |
| 2 | 0.29 | 0.342 |
| 3 | 0.39 | 0.386 |
| 4 | 0.57 | 0.568 |
| 5 | 0.38 | 0.477 |
| 6 | 0.25 | 0.309 |
| 7 | 0.34 | 0.333 |
| 8 | 0.35 | 0.379 |
| 9 | 0.4 | 0.496 |
| 10 | 0.62 | 0.596 |
| 11 | 0.67 | 0.692 |
| 12 | 0.5 | 0.59 |
| 13 | 0.61 | 0.65 |
| 14 | 0.46 | 0.36 |
| 15 | 0.69 | 0.718 |
| 16 | 0.45 | 0.497 |
| 17 | 0.51 | 0.519 |
| 18 | 0.71 | 0.74 |
| 19 | 0.49 | 0.53 |
| 20 | 0.49 | 0.511 |
| 21 | 0.59 | 0.597 |
| 22 | 0.56 | 0.553 |
| 23 | 0.49 | 0.581 |
| 24 | 0.49 | 0.596 |
| 25 | 0.75 | 0.787 |
| 26 | 0.55 | 0.603 |
| 27 | 0.53 | 0.543 |
| 28 | 0.48 | 0.578 |
| 29 | 0.48 | 0.555 |

| | | |
|---|---|---|
| 30 | 0.58 | 0.652 |
| 31 | 0.66 | 0.665 |
| 32 | 0.7 | 0.718 |
| 33 | 0.43 | 0.424 |
| 34 | 0.63 | 0.648 |
| 35 | 0.56 | 0.623 |
| 36 | 0.33 | 0.442 |
| 37 | 0.22 | 0.272 |
| 38 | 0.35 | 0.387 |
| 39 | 0.31 | 0.357 |
| 40 | 0.4 | 0.48 |
| 41 | 0.47 | 0.455 |
| 42 | 0.37 | 0.43 |
| 43 | 0.38 | 0.432 |
| 44 | 0.44 | 0.52 |
| 45 | 0.4 | 0.386 |
| 46 | 0.46 | 0.481 |
| 47 | 0.57 | 0.603 |
| 48 | 0.67 | 0.737 |
| 49 | 0.43 | 0.464 |
| 50 | 0.6 | 0.496 |
| 51 | 0.36 | 0.363 |
| 52 | 0.54 | 0.582 |
| 53 | 0.61 | 0.626 |
| 54 | 0.52 | 0.629 |
| 55 | 0.54 | 0.6 |

**Supplemental Table 4: Quantification of asymmetric and symmetric pairs in H3 and H3T3A intestines.**

| Intestine # | Asymmetric ISCs | Symmetric ISCs | Total | Percentage Asymmetric | Percentage Symmetric |
|---|---|---|---|---|---|
| **H3** | | | | | |
| 1 | 130 | 31 | 161 | 80.7 | 19.3 |
| 2 | 164 | 28 | 192 | 85.4 | 14.6 |
| 3 | 418 | 142 | 560 | 74.6 | 25.4 |
| 4 | 460 | 119 | 579 | 79.4 | 20.6 |
| 5 | 428 | 127 | 555 | 77.1 | 22.9 |
| 6 | 235 | 59 | 294 | 79.9 | 20.1 |
| **H3T3A** | | | | | |
| 1 | 207 | 209 | 416 | 49.8 | 50.2 |
| 2 | 79 | 112 | 192 | 41.1 | 58.9 |
| 3 | 297 | 220 | 517 | 57.4 | 42.6 |
| 4 | 284 | 205 | 489 | 58.1 | 41.9 |
| 5 | 360 | 374 | 734 | 49.0 | 51.0 |
| 6 | 194 | 204 | 399 | 48.9 | 51.1 |

**Supplemental Table 5: Cluster analysis of ISCs for H3 and H3T3A intestines.**

| Intestine # | 1 ISC | 2 ISCs | 3+ ISCs | Total ISCs | Total Cluster | % 1 ISC | % 2 ISCs | % 3+ ISCs | % Single ISC/ Total |
|---|---|---|---|---|---|---|---|---|---|
| **H3** | | | | | | | | | |
| 1 | 302 | 40 | 0 | 382 | 342 | 88.3 | 11.7 | 0 | 79.1 |
| 2 | 410 | 69 | 4 | 560 | 483 | 84.9 | 14.3 | 0.8 | 73.2 |
| 3 | 280 | 38 | 2 | 362 | 320 | 87.5 | 11.9 | 0.6 | 77.3 |
| 4 | 304 | 33 | 0 | 370 | 337 | 90.2 | 9.8 | 0 | 82.2 |
| **H3T3A** | | | | | | | | | |
| 1 | 123 | 57 | 7 | 259 | 187 | 65.8 | 30.5 | 3.7 | 47.5 |
| 2 | 495 | 218 | 28 | 1019 | 741 | 66.8 | 29.4 | 3.8 | 48.6 |
| 3 | 264 | 104 | 17 | 526 | 385 | 68.6 | 27.0 | 4.4 | 50.2 |
| 4 | 81 | 36 | 3 | 162 | 120 | 67.5 | 30.0 | 2.5 | 50.0 |
| 5 | 311 | 110 | 8 | 556 | 429 | 72.5 | 25.6 | 1.9 | 55.9 |
